# Glutathione S-transferase: A Candidate Gene for Berry Color in Muscadine Grapes (*Vitis rotundifolia*)

**DOI:** 10.1101/2020.07.14.202903

**Authors:** Aruna Varanasi, Margaret Worthington, Lacy Nelson, Autumn Brown, Renee Threlfall, Luke Howard, John R. Clark

**Affiliations:** Department of Horticulture, University of Arkansas, Fayetteville, AR, 72701; Department of Food Science, University of Arkansas, Fayetteville, AR, 72704

**Keywords:** glutathione S-transferase, anthocyanin, *Vitis*, berry color, muscadine grapes

## Abstract

Muscadine grapes (*Vitis rotundifolia* Michx.) are a specialty crop cultivated in the southern United States. Muscadines (2n=40) belong to the *Muscadinia* subgenus of *Vitis,* while all other cultivated grape species belong to the subgenus *Euvitis* (2n=38). The berry color locus in muscadines has been mapped to a 0.8 Mbp region syntenic with chromosome 4 of *V. vinifera*. In this study, we identified glutathione S-transferase4 (*GST4*) as a likely candidate gene for anthocyanin transport within the berry color locus. PCR and KASP genotyping identified a single intragenic SNP (C/T) marker corresponding to a proline to leucine mutation within the muscadine *GST4* (*VrGST4*) that differentiated black (CC and CT) from bronze (TT) muscadines in 65 breeding selections, 14 cultivars, and 320 progeny from two mapping populations. Anthocyanin profiling on a subset of the progeny indicated a dominant *VrGST4* action, with no allele dosage effect on total anthocyanin content or composition of individual anthocyanins. Proanthocyanidin content was similar in the seeds of both black and bronze genotypes, and seeds had much higher *VrGST3* expression and lower *VrGST4* expression than skins. *VrGST4* expression was higher in post-veraison berries of black muscadines compared to pre-veraison berries, but no changes in gene expression in pre- and post-veraison berries were observed in the bronze muscadine cultivar. *VrMybA1* expression was higher in post-veraison berries of both black and bronze muscadines. These results suggest that berry pigmentation in muscadines is regulated by a mechanism distinct from the *MybA* gene cluster that is responsible for berry color variation in *V. vinifera*.

## Introduction

Muscadine grapes (*Vitis rotundifolia* Michx.) are a specialty fruit crop native to the southeastern United States. Wild and cultivated muscadine grapes are commonly found from Delaware to central Florida and from the Atlantic coast to eastern Texas (Lane 1997). The genus *Vitis* is divided into two subgenera, *Euvitis* Planch. (bunch grapes) and *Muscadinia* Planch. All bunch grape species, such as the European wine and table grape (*V. vinifera*) and the American ‘Concord’ grape (*V. labrusca*), belong to *Euvitis*, whereas *Muscadinia* consists of only three species, *V. munsoniana* Simpson ex Munson, *V. popenoei* Fennell, and muscadine grapes (Wen 2007). Of the three species in subgenus *Muscadinia*, only muscadine grapes have commercial value (Brizicky 1965; Olien 1990). Muscadine grapes are tolerant to several pests and diseases such as grape phylloxera (*Daktulosphaira vitifoliae* Fitch) (Firoozabady and Olmo 1982), Pierce’s disease (*Xylella fastidiosa*) (Olmo 1971), and other major fungal pathogens (Staudt 1997; Merdinoglu *et al.* 2003) that cause extensive losses in *V. vinifera*.

Muscadine grapes possess distinct genetic and morphological characteristics that differentiate them from bunch grape species in subgenus *Euvitis*. Muscadine vines have unbranched tendrils and produce smaller clusters of berries that shatter at maturity. Berries are larger with thick skins and large bitter seeds (Morris and Brady 2004; Striegler *et al.* 2005). They also possess unique foxy and candy-like aroma distinct from bunch grape species (Baek *et al.* 1997). Muscadines and other *Muscadinia* grapes (2x=2n=40) also differ in the number of somatic chromosomes from bunch grapes (2x=2n=38). They are considered a source of genetic variation and disease resistance for the *Euvitis* species but are not commonly used for hybridization with bunch grapes due to the difference in the chromosomes (Olmo 1986) and graft-incompatibility (Olien 1990). However, muscadine cultivation has gained importance in recent years for fresh fruit, wine, and juice production. Furthermore, the dried pomace (left over fruit after wine and juice processing) is being used as a functional food due to its high nutraceutical content (Conner 2009; Vashisth *et al.* 2011).

Muscadine breeding is complex since it requires simultaneous selection of several important traits. The most common challenge for muscadine breeding programs is to increase fruit quality while retaining disease resistance and vine vigor. Breeding efforts are currently focused on selection for seedlessness, hermaphroditism, and stable berry color during processing (Lu *et al.* 1993; Conner and MacLean 2013; Xu *et al.* 2014; Conner *et al.* 2017; Lewter *et al.* 2019). Berry color of muscadine grapes is very important especially for wine and juice industry as poor color stability can result in reduced quality of processed products (Morris and Brady 2004; Vasanthaiah *et al.* 2011). While muscadines cover a spectrum of berry shades, there are two primary color types; black (very dark purple) or bronze (greenish yellow) (Conner and MacLean 2013). Black-fruited muscadines are more common in the wild, though bronze muscadines are also occasionally found and bronze berry color is recessively inherited (Stucky 1919). Both black and bronze muscadine cultivars are available for the fresh market and processing industries.

The color of muscadine berries is determined by the quantity and composition of anthocyanins, which are accumulated in the skins. Quality and color stability of muscadine juice and wine are affected by the total anthocyanin content, anthocyanin composition, and intra-molecular copigmentation (Sims and Bates 1994; Talcott *et al.* 2003). Six types of anthocyanidins (malvidin, peonidin, pelargonidin, petunidin, cyanidin, and delphinidin) have been identified in *V. vinifera* and muscadine grapes (He *et al.* 2010; Conner and MacLean 2013). While the anthocyanins in bunch grape species are acylated 3-O-monoglucosides, among which malvidin is more common (Janvary *et al.* 2009), muscadine grapes have non-acylated 3,5-O-diglucosidic anthocyanins with delphinidin being the predominant type (Ballinger *et al.* 1974; Sandhu and Gu 2010). Conner and McLean (2013) examined anthocyanin content and composition from the berry skin extracts of 22 *V. rotundifolia* cultivars and *Muscadinia* germplasm and found that anthocyanin content ranged from less than 100 μg.g^−1^ in bronze muscadines to over 5000 μg.g^−1^ in the highly pigmented black muscadines.

Anthocyanins are synthesized by the flavonoid biosynthetic pathway, which is one of the best studied pathways in plants (He *et al.* 2010; Tanaka and Sasaki 2008). Anthocyanin biosynthesis in plants occurs within the cytosol and the end products are transported to be deposited in the vacuoles. Regulation of genes involved in anthocyanin biosynthesis has been studied in several plant species (Cone *et al.* 1986; Quattrocchio *et al.* 1993; Holton and Cornish 1995; Zhao *et al.* 2015). While the coordinated expression of different structural genes appears to be regulated by the transcription factors, MYC and MYB proteins (Kobayashi *et al.* 2002; Deluc *et al.* 2006), their transport to the final site of storage, which is a critical process for pigmentation, is facilitated by several mechanisms including glutathione *S*-transferase (GST)- mediated transport (Dixon *et al.* 2010). GSTs comprise a large multigenic family in plants with diverse functions in the cell. They are soluble membrane-associated dimers known to function enzymatically in cellular detoxification through conjugation of xenobiotic substrates with glutathione (GSH) (Dixon and Edwards 2005). In addition, GSTs function in a non-catalytic role by acting as carrier proteins (ligandins) for shuttling of several endogenous compounds including vacuolar sequestration of anthocyanins (Dixon *et al.* 2010). GSTs are known to play a major role in anthocyanin transport in several plants such as *ZmBz2* from maize (*Zea mays*) (Marrs *et al.* 1995), *PhAn9* from petunia (*Petunia hybrida*) (Mueller *et al.* 2000), *AtTT19* from *Arabidopsis thaliana* (Sun *et al.* 2012), and *VvGST4* from *V. vinifera* (Conn *et al.* 2008; Pérez-Díaz *et al.* 2016).

Genes controlling berry color have been well characterized in *V. vinifera*. The color locus in *V. vinifera* was mapped to a 5 Mbp region on chromosome 2 and was associated with a single *MybA* gene cluster that accounted for 84% of the color variation (Fournier-Level *et al.* 2009; Myles *et al.* 2011). This variation in berry color is the result of additive effects from alleles of three MYB-type transcription factor genes, *VvMybA1*, *VvMybA2*, and *VvMybA3*, within the single *MybA* gene cluster. While *VvMybA1* and *VvMybA2* were demonstrated to be functionally involved in berry pigmentation (Kobayashi *et al.* 2002; Walker *et al.* 2007), the association of *VvMybA3* was determined purely through statistics-based approach due to the strong LD detected with *VvMybA2* (Fournier-Level *et al.* 2009). Five polymorphisms (one retrotransposon, three SNPs, and one 2-bp In/Del) within these *MybA* genes caused structural changes in the *MybA* promoters and MYB proteins that resulted in the quantitative variation of berry color in *V. vinifera* (This *et al.* 2007; Fournier-Level *et al.* 2009). White color of *V. vinifera* berries is inherited as a recessive trait and has been linked to the presence of a single gypsy-type retrotransposon, *Gret1*, in homozygous condition in the promoter of *VvMybA1* (Kobayashi *et al.* 2004; This *et al.* 2007; Fournier-Level *et al.* 2009). Although the *Gret1* transposon in *VvMybA1* is a major determinant for generating color variation, the three SNPs detected in the exon of *VvMybA1* and the single In/Del identified in the exon of *VvMbA2* also contributed significantly (23% of the variance) to the quantitative variation of anthocyanin content in *V. vinifera* (Fournier-Level *et al.* 2009).

So far, limited research has been conducted on the genetic control of berry color and its association with anthocyanin content in *V. rotundifolia*. Recent technological advances in *V. vinifera* genomics (Venturini *et al.* 2013; Fennell *et al.* 2015; Chin *et al.* 2016; Yang *et al.* 2016) and the close relationship between *V. vinifera* and *V. rotundifolia* genomes have enabled the application of genomic resources and tools from *V. vinifera* to advance molecular genetic analysis in *V. rotundifolia* (Lewter *et al.* 2019). Using two biparental F1 mapping populations segregating for flower sex and berry color with ‘Black Beauty’ or ‘Supreme’ being the female parent and ‘Nesbitt’ as the common male parent, Lewter *et al.* (2019) developed the first saturated genotyping by sequencing-based linkage maps in muscadine grape. All three parents were heterozygous for black phenotype and the progeny in both populations segregated at a 3:1 ratio for black to bronze berry color. The dense linkage maps were each composed of 20 linkage groups (LGs), with over 1200 markers in the ‘Black Beauty’ x ‘Nesbitt’ population and over 2000 markers in ‘Supreme’ x ‘Nesbitt’ population. A high degree of colinearity was observed between these genetic maps and the physical map of *V. vinifera* except for LGs 7 and 20, suggesting a highly conserved genome structure between the two species. The markers from LGs 7 and 20 of *V. rotundifolia* co-localized to chromosome 7 in *V. vinifera* suggesting a possible split of chromosome 7 in *V. vinifera* or a fusion of the two chromosomes from *V. rotundifolia* during *Vitis* evolution (Lewter *et al.* 2019).

The berry color locus mapped to a region corresponding to 11.1 – 11.9 Mbp on chromosome 4 on the physical map of *V. vinifera* in both muscadine populations (Lewter et al., 2019). This is an interesting finding, as the berry color locus in *V. vinifera* was mapped to chromosome 2 where *MybA* genes were identified as candidate genes (Fournier-Level *et al.* 2009) and suggests that other genes in the anthocyanin biosynthesis pathway possibly determine berry color in *V. rotundifolia*. Therefore, the objective of this study was to identify potential candidate genes within the 0.8 Mbp mapped locus on chromosome 4 of *V. vinifera* and validate their association with berry color variation in *V. rotundifolia*.

## Materials and Methods

### Plant material

Four muscadine cultivars used as parents or grandparents for the two biparental F_1_ mapping populations (‘Black Beauty’ x ‘Nesbitt’ and ‘Supreme’ x ‘Nesbitt’) developed for mapping the color locus (Lewter *et al.* 2019) were used in this study to search for potential candidate genes. While the three parents, ‘Black Beauty’, ‘Supreme’, and ‘Nesbitt’, are heterozygous for black berry color, ‘Fry’ is the recessive bronze-fruited cultivar found prominently in the pedigree of all three black-fruited parents (Goldy and Nesbitt 1985; Clark 1997; Conner 2013; Lewter *et al.* 2019). Plant materials (young leaves and berries) for all the muscadine samples used in this study were collected from the University of Arkansas System Division of Agriculture Fruit Research Station (FRS) in Clarksville, AR.

### Identification of a candidate gene in the V. vinifera reference genomea

The annotated genes within the 11.1 to 11.9 Mbp interval on chromosome 4 of the *V. vinifera* reference genome were explored to identify genes associated with anthocyanin biosynthesis and compared with known sequences in the NCBI database using BLAST. The *GST4* candidate gene from the four muscadine cultivars was amplified using primers designed from the *VvGST4* sequence. For initial PCR, genomic DNA was used to amplify the *GST4* sequence using forward primer 5’ ATATCAAGCAGCGAGCTCCA 3’ and reverse primer 5’ CCTCTTGGGAAAAAGCTTGG 3’. To isolate the full-length *GST4* sequence, forward primer 5’ ATATCAAGCAGCGAGCTCCA 3’ and reverse primer 5’ GGTGGAAGATGGTGATGAAGGT 3’ were used on cDNA. Sequences were compared with known *GST4* sequences from related species using BLAST. A Kompetitive Allele Specific PCR (KASP) assay for a single SNP identified in the *VrGST4* sequence was designed using two forward primers (5’ CTCGCTGATCTGAGTCATCTTCT 3’ and 5’ CTCGCTGATCTGAGTCATCTTCC 3’) and one common reverse primer (5’ CCAGCTTCCTTCACCAAGTTTCTGAT 3’). Genotyping was performed on 65 breeding selections, 14 cultivars, and 320 progeny from the two mapping populations (‘Black Beauty’ x ‘Nesbitt’ and ‘Supreme’ x ‘Nesbitt’). Deduced protein sequences of VrGST4 from the four muscadine cultivars used for candidate gene isolation were aligned with GSTs associated with anthocyanin pathway from diverse plant species using ClustalW (Thompson *et al.* 1994). Based on the alignment, a phylogenetic tree was generated using PhyML v20160115 (https://www.genome.jp/tools/ete/).

### Berry color and anthocyanin profiling

Berry samples for anthocyanin profiling were harvested from the four muscadine cultivars (‘Black Beauty, ‘Fry’, ‘Nesbitt’, and ‘Supreme’) and a subset of progeny from the two biparental mapping populations (‘Black Beauty’ x ‘Nesbitt’ and ‘Supreme’ x ‘Nesbitt’). Forty-eight progeny, 16 from each genotype class (CC, CT, or TT), were randomly selected from each population for anthocyanin profiling. Skin color at the equator was measured using a CR400 colorimeter (Konica Minolta, Ramsey, NJ) for five berries from each genotype. Afterwards, the skins and seeds from each sample were separated and used for analysis of total anthocyanins and proanthocyanidins, respectively. Phenolics from the five berry skins from each sample were extracted following Cho *et al.* (2004) and total anthocyanin content was determined using the pH differential method (Giusti and Wrolstad 2001). A subset of five randomly selected samples from each genotype class in each population were evaluated for individual anthocyanins using high performance liquid chromatography (HPLC) (Cho *et al.* 2004). The seeds from each sample were ground into a fine powder and total proanthocyanidins present in the phenolic extract were measured using the DMAC assay following Prior *et al.* (2010).

#### Gene expression analysis

Transcript levels for *VrGST4* were determined from pre-veraison and post-veraison berry skins for the four muscadine cultivars by quantitative PCR (qPCR). All qPCR reactions were performed in triplicate with two biological replicates for each sample. *VrGST4* expression was normalized using *VvUbiquitin* as the reference (Bogs 2005).

Seeds and berry skins from post-veraison berries were collected from two bronze (AM-53 and ‘Doreen’) and two black (AM-83 and ‘Nesbitt’) muscadine genotypes to study the expression of *VrGST3*, *VrGST4*, and *VrMybA1* genes. RNA was isolated from berry skins using Spectrum™ plant total RNA kit (Sigma-Aldrich, St. Louis, MO) according to the manufacturer’s instructions and from the seeds as described by Reid *et al.* (2006). Total RNA from berry skins and seeds was used for the synthesis of first strand cDNA using iScript cDNA synthesis kit (Bio-Rad Laboratories Inc., Hercules, CA) and diluted 1:10 for qPCR reactions. Gene transcript levels were measured by qPCR using a CFX96 Touch Real-Time PCR Detection System (Bio-Rad Laboratories Inc., Hercules, CA). All qPCR reactions were carried out in triplicate and expression for the three genes was normalized against *VvUbiquitin* (Bogs 2005).

### Data availability

File S1 contains a list with detailed information for all the supplementary tables and figures. More detailed information about the materials and methods used in this study can be found in File S2. Sequencing data from this project has been deposited in NCBI GenBank with the following accession numbers; VrGST4-‘Fry’: MT678460; VrGST4-‘Supreme’: MT678461; VrGST4-‘Black Beauty’: MT678462; VrGST4-‘Nesbitt’: MT678463. Other data generated and analyzed during this study are included in this published article and its supplementary information files.

## Results

### Identification and characterization of VrGST4

The *V. vinifera* genome was used as the reference to identify potential candidate gene(s) for muscadine berry color variation within the 0.8 Mbp locus identified by Lewter *et al.* (2019). Twenty-one genes annotated with known or putative function were identified between 11.1 to 11.9 Mbp on chromosome 4 of the 12X.0 version of the PN40024 *V. vinifera* reference genome. Among those, only one full-length gene, annotated as *VvGST4*, was associated with the anthocyanin biosynthetic pathway (Alfenito *et al.* 1998; Conn *et al.* 2008). Initially, a 395 bp genomic DNA fragment was amplified from the leaf tissues of black and bronze-fruited muscadines using the *VvGST4* sequence information (Figure S1). A BLASTn search of the 395 bp product was performed to compare it with known GST4 sequences from other species. The amplified 395 bp sequence from muscadine showed an identity of 98% with *VvGST4* and *V. amurensis* (*VaGST4*; Genbank Accession No. FJ645770.1) sequences suggesting that *V. rotundifolia* GST4, designated as *VrGST4*, is the likely candidate gene for berry color variation in muscadine (Table S1).

A full-length *VrGST4* cDNA of 642 bp was then amplified from berry skins of a bronze-fruited muscadine cultivar, ‘Fry’, and three black-fruited muscadine cultivars, ‘Supreme’, ‘Black Beauty’, and ‘Nesbitt’. Five SNPs were detected among the four *VrGST4* sequences, with only one nonsynonymous polymorphism (C/T) identified towards the 3’ end of the gene at position 512 (Figure 1A). Figure 1B shows the chromatograms of the nonsynonymous SNP region in the bronze and black muscadine sequences with CCG codon identified in the black muscadines and CTG codon in the bronze muscadine. The overlapping signals for C or T at position 512 in the three black muscadines indicates the presence of both CCG and CTG alleles in these cultivars.

**Figure 1.**
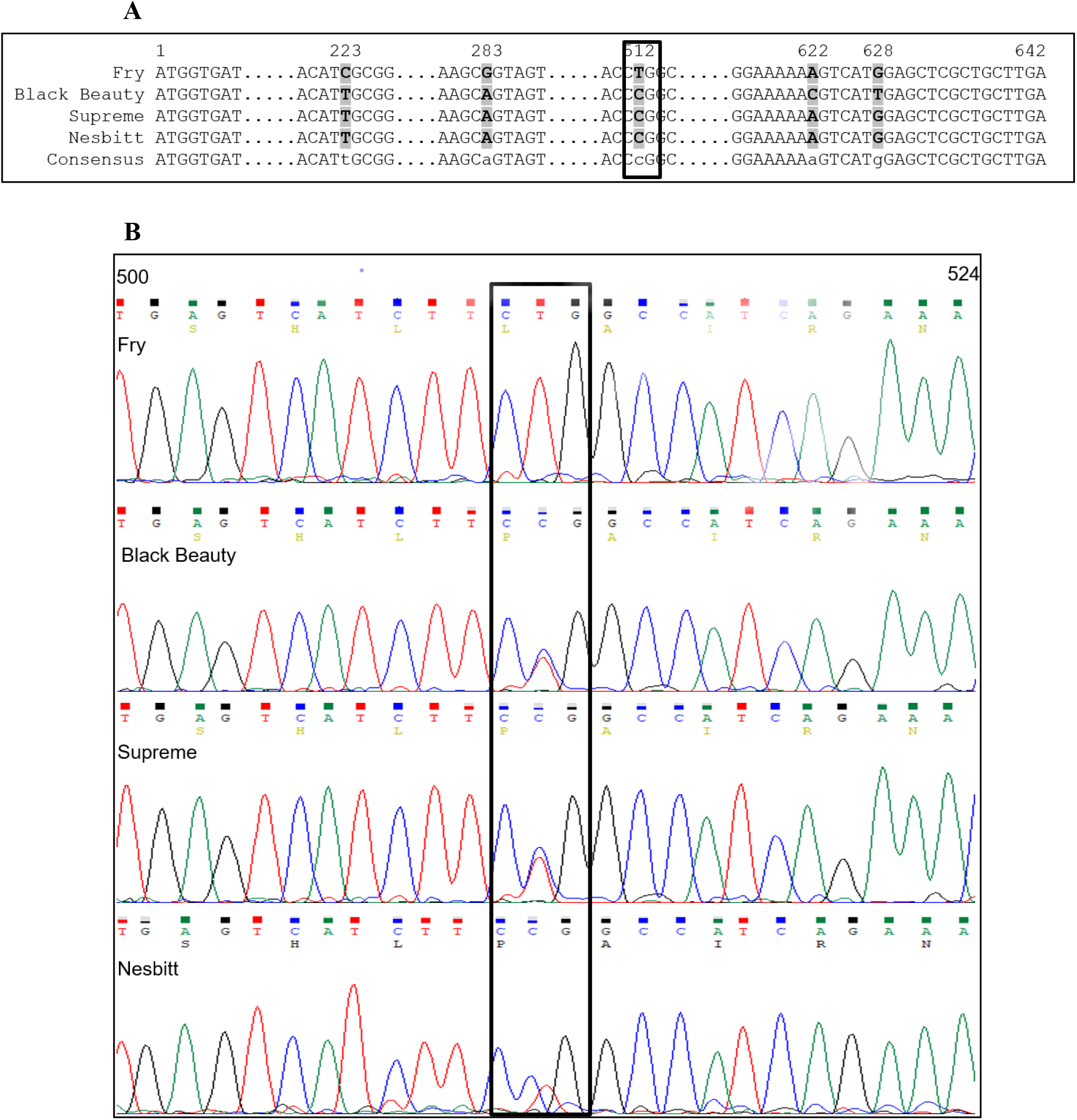
*VrGST4* cDNA analysis. (A) cDNA alignment of bronze (‘Fry’) and black (‘Black Beauty’, ‘Supreme’, ‘Nesbitt’) muscadines. Dots represent the sequences with no polymorphisms. SNPs are represented by grey highlights and the nonsynonymous SNP is represented by a black box. (B) Chromatograms of the nonsynonymous SNP region in the bronze and black sequences with the CTG/CCG codon highlighted in the black box. Peaks represent the corresponding nucleotide in the sequence. Numbers above the sequence indicate nucleotide positions.

The deduced protein from the full-length *VrGST4* sequence of both bronze and black-fruited muscadines had 213 amino acids with a predicted molecular weight of 24.2 kDa in all four muscadine cultivars. A BLASTp search of the VrGST4 protein sequences was performed to compare the sequence similarity with known GST4 proteins from other species (Figure 2). VrGST4 protein had 98, 65, and 56% sequence identity with VvGST4 (NP_001267869) from *V. vinifera*, PhAN9 (CAA68993) from *Petunia hybrida*, and AtTT19 (AED92398) from *Arabidopsis thaliana*, respectively. Protein alignment indicated that the C/T SNP identified in *VrGST4* cDNA corresponded to a proline to leucine mutation, thereby confirming the presence of an intragenic SNP marker within the candidate VrGST4 gene. Leucine was present at position 171 in bronze muscadine, whereas proline was present at this position in VvGST4, PhAN9, AtTT19, and VrGST4 proteins of black muscadines.

**Figure 2.**
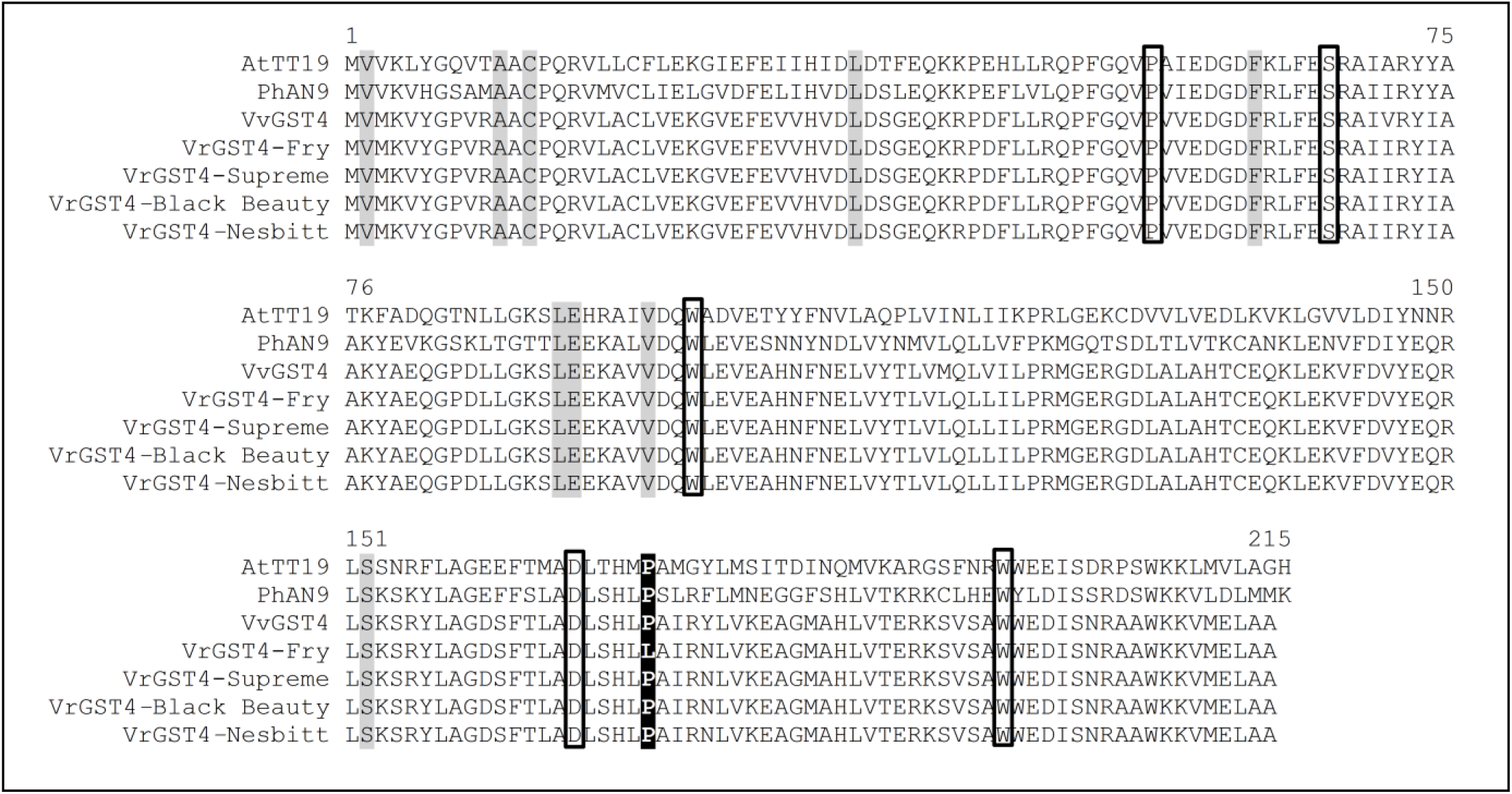
VrGST4 protein alignment. Comparison of VrGST4 from post-veraison berry skins of ‘Fry’, ‘Supreme’, ‘Black Beauty’, and ‘Nesbitt’ cultivars with anthocyanin-related GST sequences from Arabidopsis (AtTT19), Petunia (PhAN9), and *V. vinifera* (VvGST4). Numbers above the sequence alignment represent amino acid positions. Black shaded region indicates the proline to leucine mutation at position 171. Grey shaded regions indicate amino acid residues specific to anthocyanin-related GSTs (Kitamura *et al.* 2012). Black boxes indicate amino acid residues suggested previously to have high homology in anthocyanin-related GSTs (Alfenito *et al.* 1998).

A phylogenetic tree derived from the deduced VrGST4 protein sequences of bronze and black muscadines, and other known GSTs (Figure 3) indicated that VrGST4 belonged to the Phi group of GST proteins. Results showed that VrGST4 from both bronze and black muscadines is closely related to VvGST4 (Conn *et al.* 2008; Perez-Dias *et al.* 2016) and VaGST4 from *V. amurensis* and grouped in the same clade as PhAN9 (Alfenito *et al.* 1998; Mueller *et al.* 2000), AtTT19 (Kitamura *et al.* 2004), and other Phi group GSTs such as CkmGST3 of *Cyclamen sp.* (Kitamura *et al.* 2012), CsGST of *Citrus sinensis* (Licciardello *et al.* 2014), and LcGST4 of *Litchi chinensis* (Hu *et al.* 2016). These Phi group GSTs were all previously reported to function as anthocyanin transporters in the flavonoid biosynthetic pathway. Other anthocyanin-related GSTs belonging to the Tau group, ZmBz2 of *Zea mays* (Marrs *et al.* 1995) and VvGST1 of *V. vinifera* (Perez-Dias *et al.* 2016), clustered separately.

**Figure 3.**
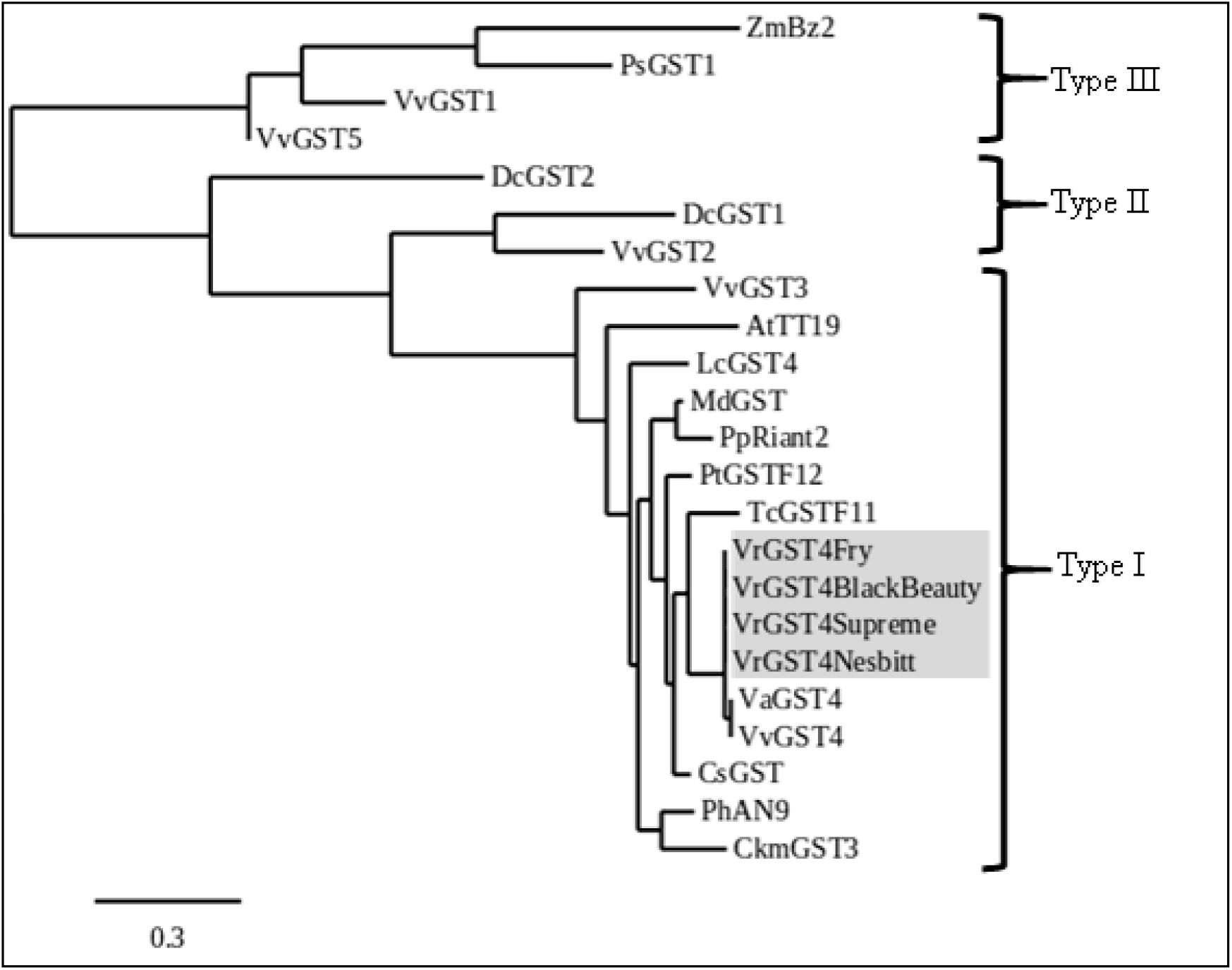
Phylogenetic tree derived from deduced protein sequences of VrGST4 (grey box) from bronze (‘Fry’) and black (‘Black Beauty’, ‘Supreme’, ‘Nesbitt’) muscadines relative to protein sequences of known GSTs from other species retrieved from GenBank. GST classification is represented by Type I, II, and III classes. At, *Arabidopsis thaliana* (AED92398); Ckm, *Cyclamen sp*. (BAM14584); Cs, *Citrus sinensis* (ABA42223); Dc, *Dianthus caryophyllus* (DcGST2, AAA51450; DcGST1, AAA72320); Lc, *Litchi chinensis* (KT946768); Md, *Malus domestica* (AEN84869); Ph, *Petunia hybrida* (CAA68993); Pp, *Prunus persica* (KT312848); Ps, *Papaver somniferum* (AAF22517); Pt, *Populus trichocarpa* (XP_006372485); Tc, *Theobroma cacao* (EOY03123); Va, *Vitis amurensis* (ACN38271); Vr, *Vitis rotundifolia* (VrGST4); Vv, *Vitis vinifera* (VvGST1, AAN85826; VvGST2, ABK81651; VvGST3, ABO64930; VvGST4, NP_001267869; VvGST5, ABL84692); Zm, *Zea mays* (AAA50245).

### Validation of VrGST4 SNP marker in diverse genotypes

In order to further validate the association of the intragenic C/T SNP with berry color in diverse muscadine genotypes, KASP genotyping assay was performed on 320 progeny from the two mapping populations, 65 breeding selections from the Arkansas Fruit Breeding Program, and 14 cultivars including the four muscadine parents selected for identification and characterization of *VrGST4* (Table S1). The results are depicted as a cluster plot (Figure 4) showing three clusters of data points representing homozygous black (CC), heterozygous black (CT), and bronze-fruited (TT) genotypes. Association of KASP marker data with berry color from all the genotypes revealed that CC and CT genotypes associated with black phenotype whereas TT genotype associated with the bronze phenotype. The marker reaction failed for 17 of 399 samples and no genotype was predicted. The KASP marker correctly predicted berry color in 376 of the 382 successful reactions. New leaf tissue was collected from the six vines with phenotype data that did not match the KASP genotype prediction and the full-length *VrGST4* fragment was amplified from those samples. In five of six cases, the new sequence data matched the berry color phenotype, indicating that the discrepancies between marker predictions and phenotypes were caused by errors in leaf collection among tightly spaced plants in the research vineyard.

**Figure 4.**
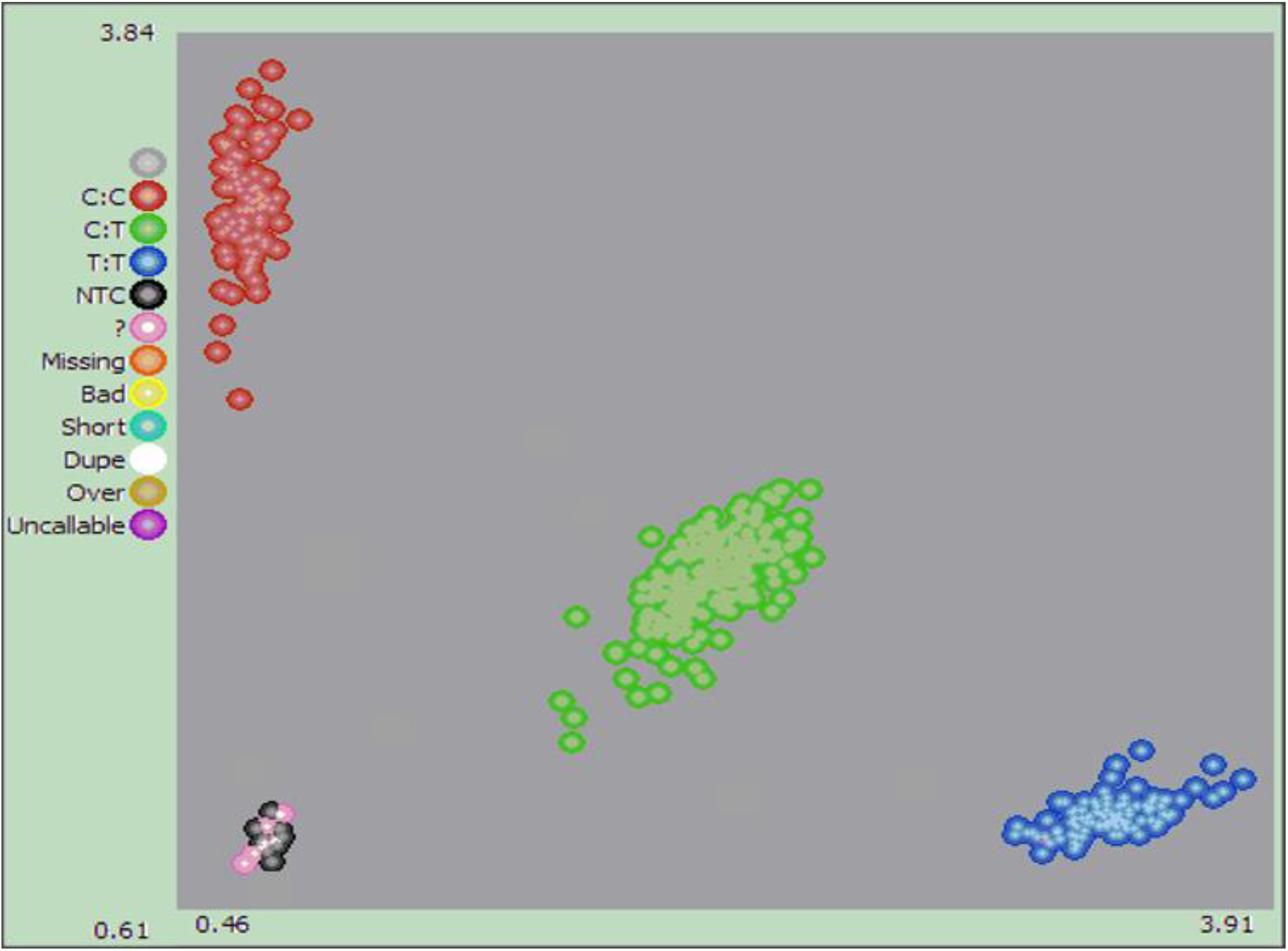
Cluster plot analysis from KASP genotyping assay using an intragenic SNP (C/T) marker of *VrGST4*. Red, green, and blue clusters represent CC, CT, and TT genotypes, respectively from diverse muscadine genotypes segregating for black (CC & CT) or bronze (TT) berry color. The pink dots represent samples that could not be determined and the black cluster represents negative controls.

### Expression of VrGST4 in pre-veraison and post-veraison berries

A qPCR assay was performed to detect the level of *VrGST4* expression in berry skins from three black muscadine cultivars and one bronze cultivar (Figure 5). *VrGST4* expression was depicted as fold expression relative to *VvUbiquitin* reference. In all three black muscadines, *VrGST4* expression was significantly higher in the post-veraison berries than the pre-veraison berries. ‘Nesbitt’ had the highest level of *VrGST4* expression in both pre-veraison (0.52) and post-veraison (1.27) berries compared to ‘Supreme’ (0.09 in pre-veraison and 0.72 in post-veraison berries) and ‘Black Beauty’ (0.11 in pre-veraison and 0.62 in post-veraison berries). However, the fold increase of *VrGST4* levels in post-veraison berries compared to pre-veraison berries was highest in ‘Supreme’ (8-fold), followed by ‘Black Beauty’ (6-fold), and ‘Nesbitt’ (2.5-fold). Interestingly, the bronze cultivar, ‘Fry’ had higher levels of *VrGST4* expression in pre-veraison (1.22) berries than all the three black muscadines. Furthermore, the post-veraison *VrGST4* expression in ‘Fry’ (1.15) was comparable to ‘Nesbitt’ but higher than both ‘Supreme’ and ‘Black Beauty’. However, unlike in the three black muscadines, there was no difference in expression level between the pre-veraison and post-veraison berries of the bronze cultivar.

**Figure 5.**
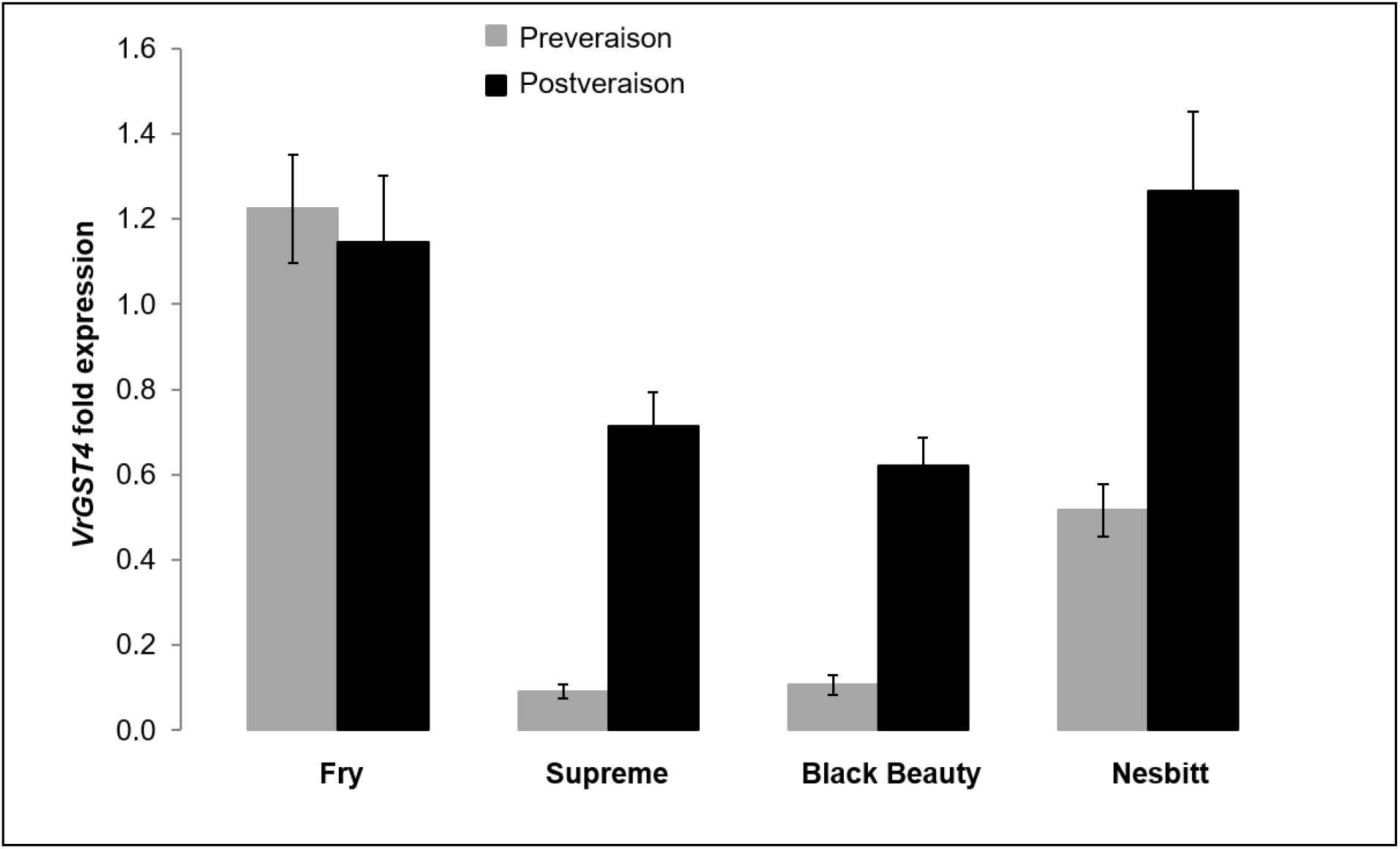
Expression of *VrGST4* from pre-veraison and post-veraison berry skins relative to *VvUbiquitin* in bronze (‘Fry’) and black (‘Supreme’, ‘Black Beauty’, ‘Nesbitt’) cultivars. Data represents means of two biological samples. Error bars represent standard error of mean.

### Berry color, total and individual anthocyanins, and total proanthocyanidins

#### Total anthocyanins and berry color

Total anthocyanin content of berry skins was estimated in a subset of progeny from the ‘Supreme’ x ‘Nesbitt’ and ‘Black Beauty’ x ‘Nesbitt’ mapping populations to determine gene action and allele dosage effects (Figure 6a). Anthocyanin content was higher in homozygote and heterozygote black genotypes of both populations compared to bronze genotypes. Average anthocyanin content across black-fruited (CC and CT) genotypes was higher in the ‘Black Beauty’ x ‘Nesbitt’ population (847.6 mg.100 g^−1^ fresh wt) compared to the ‘Supreme’ x ‘Nesbitt’ population (266.0 mg.100 g^−1^ fresh wt). The total anthocyanin content was similar in CC and CT genotypes from both populations. While in the ‘Supreme’ x ‘Nesbitt’ population, CC and CT genotypes had an average of 263.8 and 265.4 mg.100^−1^ g fresh weight, respectively, the CC and CT genotypes of ‘Black Beauty’ x ‘Nesbitt’ population averaged 890.2 and 883.1 mg.100 g^−1^ fresh weight, respectively. The TT genotypes averaged 9.4 and 18.6 mg.100 g^−1^ fresh weight in ‘Supreme’ x ‘Nesbitt’ and ‘Black Beauty’ x ‘Nesbitt’ populations, respectively. These results imply dominant gene action for *VrGST4*.

**Figure 6.**
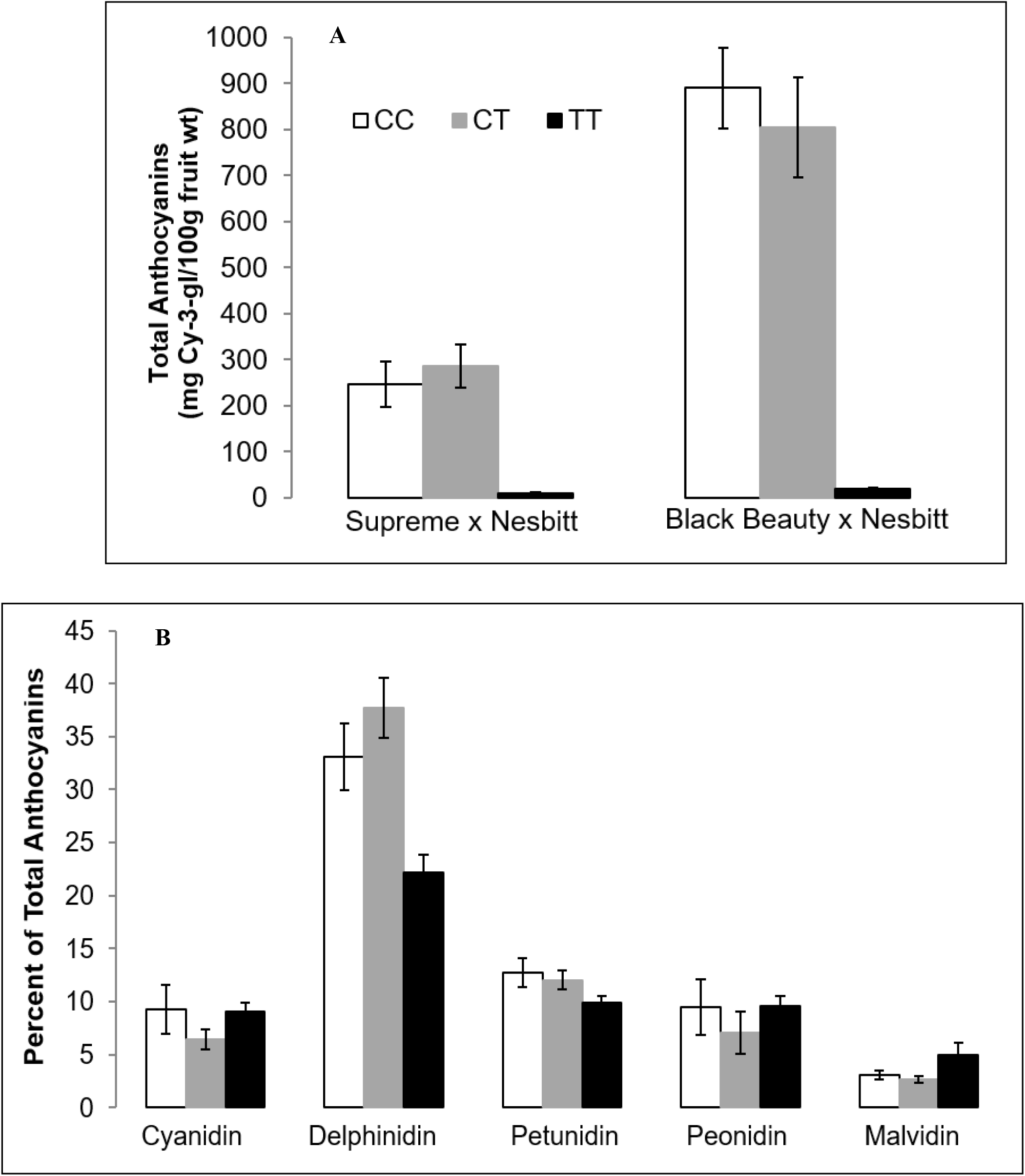
Anthocyanin profiling of post-veraison berry skins from black (CC & CT) and bronze (TT) muscadine genotypes from the ‘Supreme’ x ‘Nesbitt’ and ‘Black Beauty’ x ‘Nesbitt’ mapping populations. (A) Total anthocyanin content from each population and (B) Individual anthocyanins represented as an average of both populations. Error bars represent standard error of means.

Berry color of the black-fruited progeny (CC or CT genotypes) from the mapping populations was measured to determine the association of berry color attributes (lightness value, hue angle, and chroma) with total anthocyanin accumulation (Figure 7). The average lightness (25.81), chroma (7.27), and hue angle (10.30°) in black-fruited progeny from the ‘Supreme’ x ‘Nesbitt’ population were comparable to those of 26.39, 7.41, and 10.44°, respectively, in the ‘Black Beauty’ x ‘Nesbitt’ population. There was no significant correlation between total anthocyanins and lightness value (*P* = 0.25), hue angle (*P* = 0.35), or chroma (*P* = 0.53) in the ‘Supreme’ x ‘Nesbitt’ population. In the ‘Black Beauty’ x ‘Nesbitt’ population, there was no significant correlation between total anthocyanins and lightness (*P* = 0.05) or hue angle (*P* = 0.20). However, total anthocyanins were negatively correlated with chroma in the ‘Black Beauty’ x ‘Nesbitt’ population (*r* = −0.64, *P* < 0.001).

**Figure 7.**
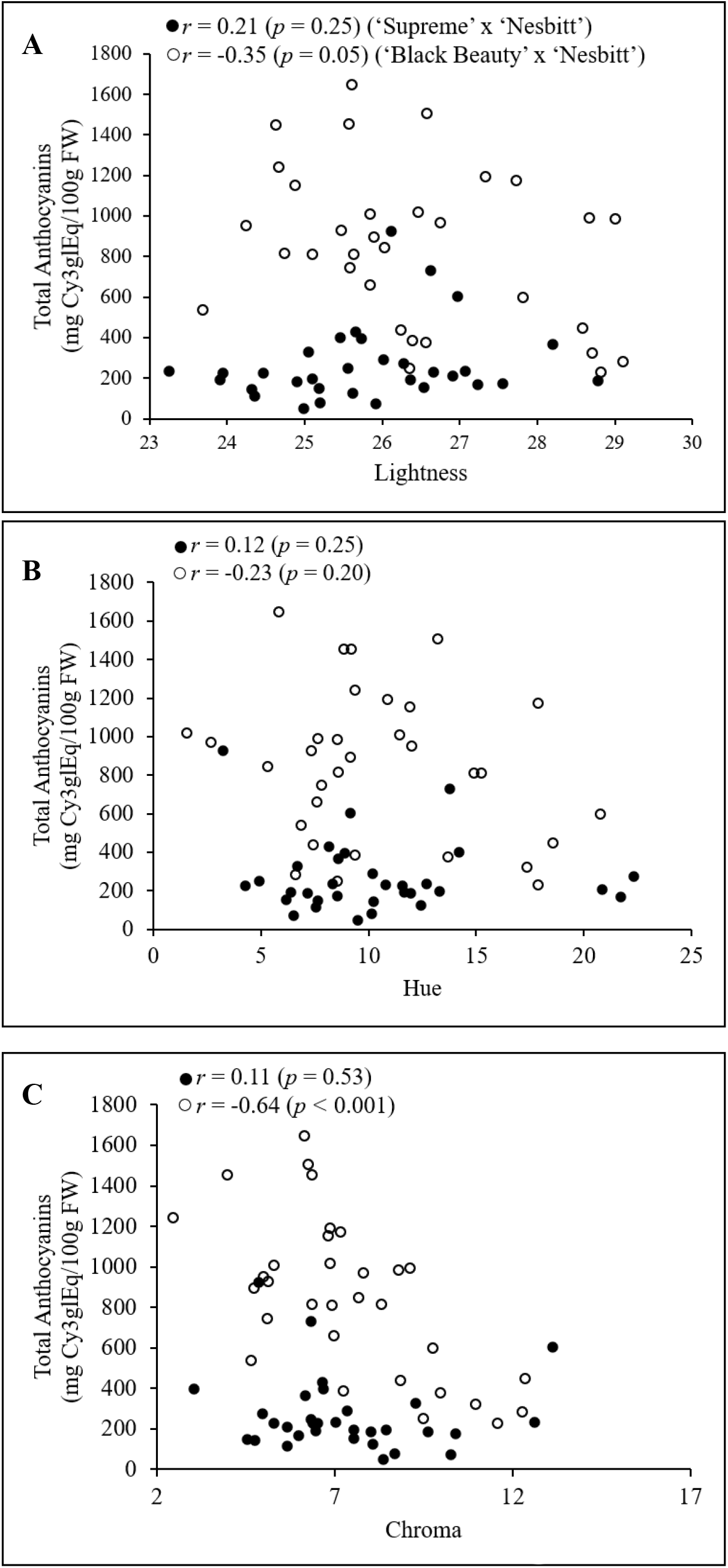
Muscadine berry color correlation (*r*) with total anthocyanin content in post-veraison berries from the two mapping populations. Color coordinates; (A) Lightness, (B) Hue, and (C) Chroma.

#### Individual anthocyanins

The composition of individual anthocyanins from a smaller subset of progeny in each mapping population was estimated using HPLC. Results are presented as percent of the total anthocyanins averaged across both mapping populations for each individual anthocyanin (Figure 6B). The percent of five individual anthocyanins, cyanidin, delphinidin, petunidin, peonidin, and malvidin, was determined from the three genotype classes (CC, CT, and TT) in both mapping populations. Delphinidin was the most abundant anthocyanin, making up 22.2 to 37.7% of total anthocyanins in all three genotype classes of both populations. This was followed by petunidin (9.9 to 12.7%), peonidin (7.1 to 9.6%), cyanidin (6.4 to 9.3%), and malvidin (2.7 to 5.0%). Petunidin, peonidin, and cyanidin content were similar among the three genotype classes. The bronze-fruited (TT) genotypes had slightly lower delphinidin content and higher malvidin content than the black genotypes. Results indicate that anthocyanin composition varied significantly between black and bronze muscadines only in delphinidin content, which is the predominant type of anthocyanin found in muscadines.

#### Total Proanthocyanidins

In addition to anthocyanin profiling, total proanthocyanidins (PAs), also known as condensed tannins, were measured from the seeds of the selected subset of progeny representing the three genotype classes (CC, CT, and TT). PA content was similar among the three genotype classes in both mapping populations (Figure 8). In the ‘Supreme’ x ‘Nesbitt’ population, CC, CT, and TT genotypes averaged 1343.2, 1103.3, and 1048.1 mg total PA 100 g^−1^ seeds, respectively. In the ‘Black Beauty’ x ‘Nesbitt’ population, CC, CT, and TT genotypes averaged 1122.0, 1170.0, and 1117.8 mg total PA 100 g^−1^ seeds. There was no significant difference in total PA content between black and bronze-fruited muscadines.

**Figure 8.**
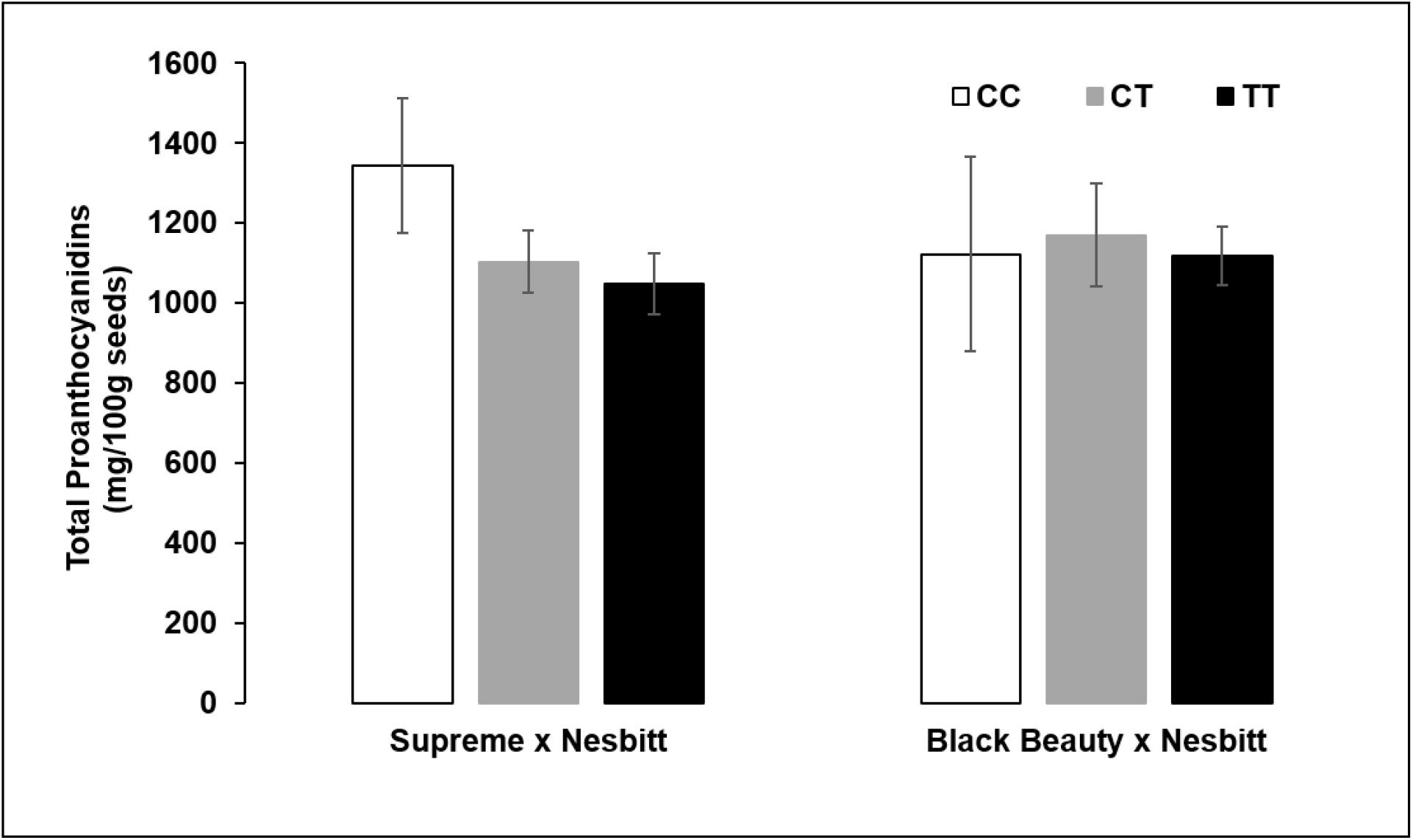
Total proanthocyanidin content from seeds of post-veraison berries from black (CC & CT) and bronze (TT) genotypes of each population. Error bars represent standard error of means.

### Differential expression of VrGST3, VrGST4, and VrMybA1 in seeds and berries

The fold expression of *VrGST3*, *VrGST4*, and *VrMybA1* from seeds and berry skins of post-veraison berries in two bronze (‘AM53’, ‘Doreen’) and two black (‘AM83’, ‘Nesbitt’) muscadine genotypes relative to *VvUbiquitin* are reported in Figures 9A, 9B, and 9C, respectively. In all four genotypes, *VrGST3* expression was higher in seeds compared to berry skins with no difference in the expression between bronze and black muscadine seeds. In contrast, as observed in Figure 5, *VrGST4* expression was higher in berry skins than in seeds. A similar trend was observed for *VrMybA1*, with higher expression observed in berry skins compared to seeds. In general, *VrGST4* expression had the greatest fold difference between seeds and skins of both bronze and black genotypes, followed by *VrGST3* and *VrMybA1* expression. *VrGST3* expression, averaged across the two bronze and two black genotypes, was 33- and 14.5-fold higher in seeds than in berry skins, respectively (Figure 9A), while average *VrGST4* expression was ~500- and ~600-fold higher in berry skins than seeds of the bronze and black genotypes, respectively (Figure 9B). On the contrary, *VrMybA1* showed only 2 to 3-fold higher expression in berry skins compared to seeds in both bronze and black genotypes (Figure 9C). Results suggest a differential expression pattern with these three genes in seeds and berry skins of bronze and black muscadines.

**Figure 9.**
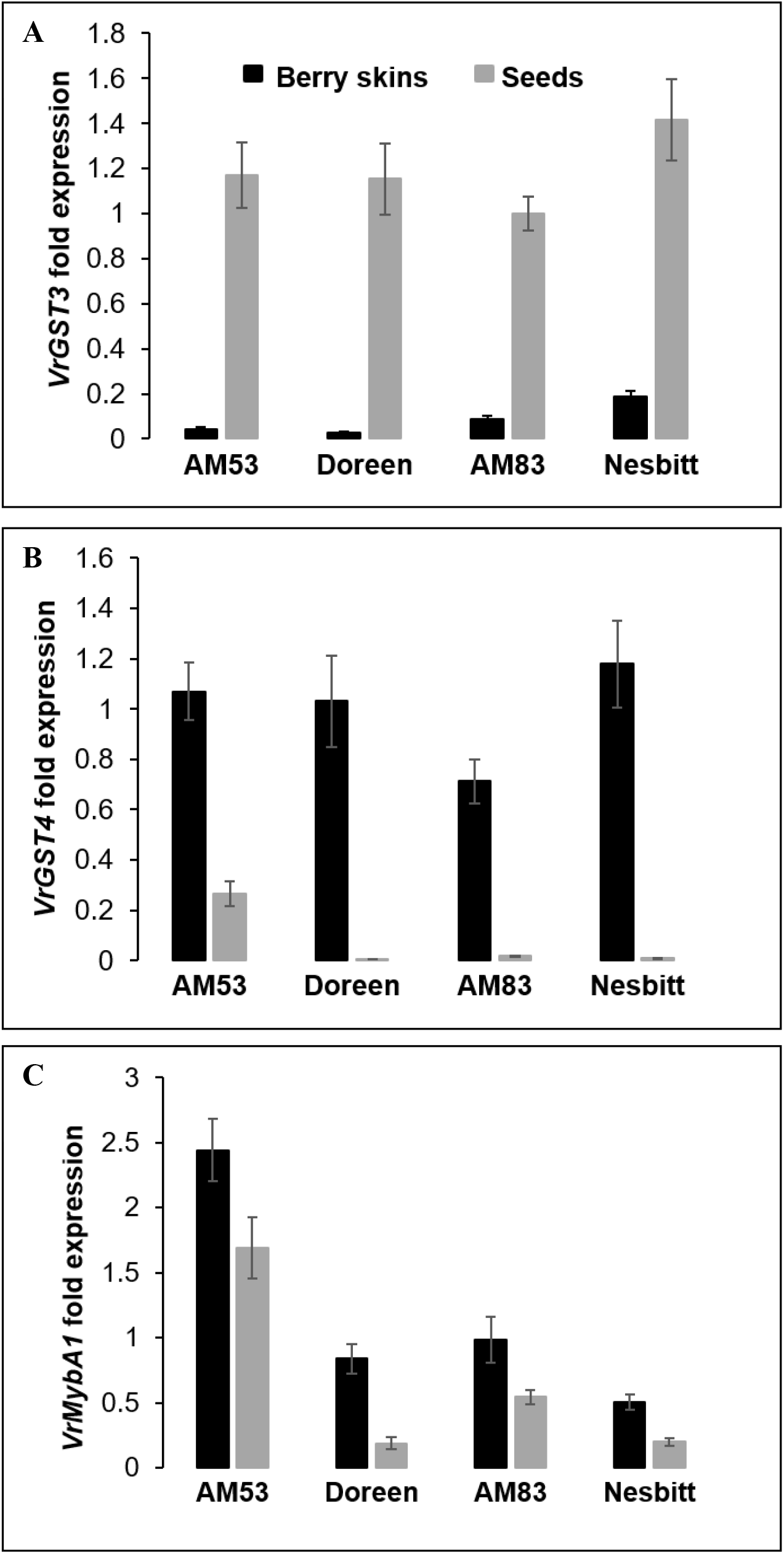
Differential gene expression analysis in bronze (‘AM53’ & ‘Doreen’) and black (‘AM83’ & ‘Nesbitt’) muscadines. Expression of (A) *VrGST3*, (B) *VrGST4*, and (C) *VrMybA1* from postveraison berry skins and seeds relative to ubiquitin. Data represents means of two biological samples. Error bars represent standard error of mean.

## Discussion

Lewter *et al.* (2019) developed the first saturated genotyping-by-sequencing-based linkage maps of muscadine grapes using two biparental mapping populations with ‘Black Beauty’ or ‘Supreme’ as the female parent and ‘Nesbitt’ as the common male parent. These dense linkage maps were used to map the muscadine berry color locus to a 0.8 Mbp region on chromosome 4 of *V. vinifera*. Variation in berry color and anthocyanin content in *V. vinifera* is controlled by a well-documented MYB gene cluster located on chromosome 2 (Kobayashi *et al.* 2004; This *et al.* 2007; Fournier-Level *et al.* 2009; Myles *et al.* 2011). Thus, the findings of Lewter *et al.* (2019) suggested that a different gene(s) within the flavonoid biosynthetic pathway was responsible for variation in berry color in muscadine grapes and that a candidate gene associated with anthocyanin accumulation and sequestration could be identified within the 11.1-11.9 Mbp interval on chromosome 4. In the present study, we demonstrated that *VrGST4*, a homolog of *VvGST4* located within the 0.8 Mbp berry color locus, is a candidate gene for berry color variation in muscadine grapes. Protein alignment of VrGST4 (Figure 2) indicated that it has high homology with VvGST4, PhAN9, and AtTT19, which have essential roles in anthocyanin transport (Perez-Dias *et al.* 2016; Mueller *et al.* 2000; Sun *et al.* 2012).

GSTs belong to a wide ubiquitous family of glutathione transferase enzymes found in bacteria, fungi, animals, and plants (Edwards *et al.* 2000). In plants, GSTs are known to perform diverse functions in the cells including defense mechanisms against pathogens, herbicide detoxification, and in pathways related to biosynthesis and detoxification of secondary metabolites such as anthocyanins (Dixon *et al.* 1998; Monticolo *et al.* 2017). GSTs are classified into six classes, Tau, Phi, Zeta, Theta, Lambda and Dhar, based on the identity of amino acid sequence. Tau and Phi are plant-specific classes due to their greater representation in terms of number of sequences (Edwards and Dixon 2005). Many GSTs have been identified in plants such as Arabidopsis (53 GSTs; Sappl *et al.* 2009), rice (59 GSTs; Soranzo *et al.* 2004), citrus (61 GSTs; Licciardello *et al.* 2014), and litchi (139 GSTs; Hu *et al.* 2016). Phylogenetic analysis based on deduced protein sequences (Figure 3) revealed that VrGST4 of both bronze and black muscadine cultivars clustered closely with GSTs from *Vitis* (VaGST and VvGST4) and formed a subclade with GSTs from other dicotyledons including AtTT19, PhAN9, and LcGST4. These closely clustered GSTs are known to perform similar function in vacuolar anthocyanin sequestration (Conn *et al.* 2008; Perez-Dias *et al.* 2016; Alfenito *et al.* 1998; Mueller *et al.* 2000; Kitamura *et al.* 2004; Hu *et al.* 2016). Furthermore, GSTs have also been classified as Type I, II or III based on their intron:exon structure (Droog 1997). According to this classification, ZmBZ2, VvGST1, and VvGST5, which are Tau class GSTs grouped as Type III class with two exons and one intron (Marrs 1996; Conn *et al.* 2008). DcGST1 and DcGST2 from carnation grouped as Type II GSTs known to have ten exons and nine introns (Marrs 1996; Sasaki *et al.* 2012), and VvGST4, PhAN9, AtTT19 clustered in the Type I class with three exons and two introns (Marrs 1996; Conn *et al.* 2008).

In this study, we initially isolated a partial genomic DNA sequence of *VrGST4* from muscadine cultivars. Hence, we could not determine the intron:exon structure directly by comparing the *VrGST4* genomic DNA with *VrGST4* cDNA. However, based on the grouping of VrGST4 protein sequence deduced from the full-length cDNA, it appears that VrGST4 from muscadine has a gene structure similar to Type I GSTs, which contain three exons and two introns. Although the Tau GSTs have a gene structure that is different from Phi GSTs, they were found to complement each other functionally in terms of anthocyanin transport. For example, ZmBZ2 and PhAN9 reciprocally complemented an9 and bz2 tissues in particle gun bombardment assays (Alfenito *et al.* 1998), even though they have only 12% amino acid identity. This functional complementation between monocot and dicot GST proteins suggests that they share a common ancestral gene before evolution into species-specific GSTs that transport different anthocyanin pigments (Alfenito *et al.* 1998).

Sequence analysis of the black and bronze muscadines isolated in our study revealed a single intragenic SNP (C/T) within the *VrGST4* cDNA corresponded to a proline to leucine mutation in the bronze muscadine (Figure 1A, Figure 2). The black muscadines used in this study were parents of the two mapping populations used for mapping the color locus (Lewter *et al.* 2019). All three black-fruited cultivars had one bronze-fruited parent and one black-fruited parent and produced progeny segregating for berry color when crossed with one another, indicating that they were all heterozygous for the berry color allele (Lewter et al. 2019). The chromatograms in Figure 1B with overlapping peaks confirm the presence of heterozygous alleles at C/T SNP in the three black muscadines, whereas the bronze muscadine had a single peak representing the homozygous recessive bronze genotype. Results from a KASP genotyping assay developed from the intragenic SNP (C/T) in *VrGST4* (Figure 4, Table S1) showed that the marker was able to distinguish between bronze (TT), heterozygote black (CT), and homozygote black (CC) genotypes and accurately predict berry color phenotype in a panel of 65 breeding selections, 14 cultivars, and 320 progeny from the two mapping populations.

Total and individual of anthocyanins were measured in a subset of progeny from each mapping population to investigate whether allele dosage (additive genetic variation) at *VrGST4* plays a significant role in determining anthocyanin content in muscadine skins. No difference in the total anthocyanin content between the homozygote and heterozygote black genotypes (Figure 6A) in either mapping populations, indicating completely dominant gene action for *VrGST4* in muscadine. In contrast, allele dosage in the MYB gene cluster plays a major role in determining anthocyanin content in *V. vinifera* and most phenotypic variation in grape anthocyanin content has been attributed to additive effects, with dominance playing a minor role (Fournier-Level et al. 2009). Our findings suggest that while the intragenic *VrGST4* KASP marker can be used to predict bronze or black berry color in breeding populations and distinguish homozygote and heterozygote black genotypes from one another, it is not useful for selecting progeny with high anthocyanin production for processing and nutraceutical industries.

Total anthocyanin content in the skins of black-fruited muscadines has previously been shown to range from less than 100 mg.100 g^−1^ to over 500 mg.100 g^−1^ (Conner and MacLean 2013). In this study, the average anthocyanin content across black-fruited (CC and CT) genotypes was over three times higher in the ‘Black Beauty’ x ‘Nesbitt’ population (847.6 mg.100 g^−1^ fresh weight) than the ‘Supreme’ x ‘Nesbitt’ population (266.0 mg.100 g^−1^ fresh weight). This discrepancy in total anthocyanin content of black-fruited progeny between the two populations could be attributed to many factors, including possible differences in ripeness on the date of harvest. The large difference between the means of the black-fruited genotypes in the two mapping populations also suggests that other loci in addition to *VrGST4* may contribute to variation in total anthocyanin content in muscadines. Further investigations are needed to determine which other loci contribute to this large range in total anthocyanin content in black muscadines.

Estimation of total anthocyanin content based on berry color is challenging, as color is not always a good predictor of nutraceutical content. Food color is a critical parameter used as a quality index and is most often described by measurements of lightness (brightness), chroma (degree of color saturation), and hue angle (color wheel with red, yellow, green, and blue at 0°, 90°, 180°, and 270°, respectively). We measured surface color characteristics in a representative sample from homozygote and heterozygote black genotypes of the two mapping populations to determine if berry color was correlated with total anthocyanin content in black muscadines (Figure 7). Although higher anthocyanin content was observed in the ‘Black Beauty’ x ‘Nesbitt’ population compared to the ‘Supreme’ x ‘Nesbitt’ population, average berry color parameters were similar in both populations. The negative correlation between chroma and total anthocyanins observed in the ‘Black Beauty’ x ‘Nesbitt’ population (Figure 6C) may be attributed to genotype-specific differences in the epicuticular wax deposition during berry development. Spinardi *et al*. (2019) observed a similar decrease in both chroma and L* values during blueberry development as the berry skin pigmentation and cuticular wax load increased resulting in darker and less vivid color of the berry peels. Overall, our results suggest that surface color characteristics are not a good predictor of total anthocyanin content in black-fruited muscadine grapes. A similar result was reported by Muthusamy *et al.* (2014), who found that color was poor indicator of beta carotene content in maize, with no significant correlation observed between color and nutraceutical content. In contrast, Palonen and Weber (2019) observed significant correlations between anthocyanin concentration and hue values and L*a*b ratios in raspberry.

Besides anthocyanin quantity, composition of individual anthocyanins also varies widely in fruits. Six anthocyanidins have been detected in *V. vinifera* grapes and muscadines, among which delphinidin and cyanidin are most commonly found in muscadines (Conner and MacLean 2013). Our results from anthocyanin composition analysis in black and bronze muscadines show that delphinidin is the predominant type of individual anthocyanidin in both black and bronze berries of the two mapping populations, which is consistent with earlier findings. Although anthocyanin quantity varied significantly between black and bronze genotypes, anthocyanin composition remained similar among the three genotype classes (Figure 7B).

Expression of *VrGST4* in pre-veraison and post-veraison berries from bronze and black-fruited muscadine cultivars were analyzed (Figure 5). The expression of *VrGST4* was consistent with anthocyanin accumulation in post-veraison berry skins of black muscadines as the pre-veraison black muscadine berries showed weak expression of *VrGST4*. However, the higher expression of *VrGST4* in both pre-veraison and post-veraison berry skins of the bronze cultivar suggests that the mutated *VrGST4* does not function in anthocyanin transport in the bronze muscadines but may be involved in the transport of alternative ligands of flavonoid origin. Further research is needed through functional complementation assays to determine the role of mutated *VrGST4* in bronze cultivars.

GSTs perform diverse functions in several pathways and can be broadly categorized into those with catalytic activity and/or ligandin activity. Catalytic activity of GSTs is used for conjugation of xenobiotic substrates with the tripeptide glutathione (GSH) and function in the detoxification of herbicides. GSTs with ligandin activity function in a non-catalytic role by acting as carrier proteins for shuttling of several endogenous compounds including vacuolar sequestration of anthocyanins (Dixon *et al.* 2002; Monticolo *et al.* 2017). GSTs from many species have been demonstrated to have a ligandin activity for the transport of anthocyanins from cytosol to vacuoles, however, their ligandin and/or catalytic activity in the transport of other flavonoids such as PAs has been studied to a lesser extent.

PAs are a group of important secondary metabolites also synthesized via the flavonoid pathway. They are generally found in seeds and contribute to the dark coloration of the seed coat. In *V. vinifera*, five GST genes (*VvGST1* to *VvGST5*) have been identified that are associated with the flavonoid biosynthetic pathway (Conn *et al.* 2008; Perez-Dias *et al.* 2016). While *VvGST1* and *VvGST4* were associated with anthocyanin transport in berries, *VvGST3* was found to play a major role in PA transport in seeds. In this study, we measured PA content in seeds of a representative sample of CC, CT, or TT genotypes from both muscadine mapping populations (Figure 8) and evaluated expression patterns of *VrGST3* and *VrGST4* in berry skins and seeds of bronze and black cultivars (Figures 9A, 9B). Expression of *VrMybA1* was also analyzed to determine its association with PA or anthocyanin transport as a regulator of *VrGST3* or *VrGST4* (Figure 9C). There was no significant difference in the PA content among the three genotype classes, indicating that bronze and black muscadines do not differ in PA content. There was differential expression of *VrGST3* and *VrGST4* in seeds and berry skins. The high expression of *VrGST3* in seeds and weak or total absence of expression in berry skins of both bronze and black muscadine genotypes, suggests that *VrGST3* is almost exclusively involved in PA transport. In contrast, the higher expression of *VrGST4* in berry skins than in seeds and equal PA content in the seeds of black and bronze-fruited muscadines implies that *VrGST4* does not have an important role in the transport and vacuolar sequestration of PAs in seeds. These results are consistent with earlier findings by Perez-Dias *et al.* (2016) in *V. vinifera*. Hu et al. (2016) also found that *LcGST4* in litchi was involved in transport of anthocyanins but not of PAs. In the case of *VrMybA1*, expression was only slightly higher in berry skins compared to seeds of both berry colors. *VrMybA1* likely regulates the expression of *VrGST4* along with other structural enzymes such as UFGT during anthocyanin biosynthesis and transport in muscadine grapes as has been previously suggested in *V. vinifera* (Fournier-Level *et al.* 2009; 2010).

## Conclusions

In this study, we isolated and characterized the candidate gene, *VrGST4*, responsible for berry color variation in muscadine grapes for the first time. We identified a non-synonymous SNP (C/T) within *VrGST4* that corresponded to a proline to leucine mutation in the bronze muscadines. A diagnostic KASP marker was developed from the intragenic SNP which co-segregated with berry color and allowed differentiation of bronze (TT), heterozygous black (CT), and homozygous black (CC) muscadines. *VrGST4* action was dominant; no allele dosage effects were observed on total anthocyanin content in black muscadines. Although PA content did not differ among black and bronze muscadine berries, expression studies indicated that different GST genes are involved in the transport of PAs (*VrGST3*) in seeds and anthocyanins (*VrGST4*) in berry skins. Further studies are needed through functional complementation assays and molecular modeling and docking to validate these functions. Our results suggest that berry color variation in muscadines is controlled by a mechanism different from that reported in *V. vinifera*. In addition to the MYB genes that regulate anthocyanin biosynthetic pathway, GST genes were identified to play a key role in the development and accumulation of pigments responsible for wine quality. These results will not only have important implications in muscadine breeding programs and the muscadine processing industry but also will provide insights in understanding the evolutionary pathways of *Vitis* species. Furthermore, this study sheds new light on the differential expression and regulation of transporter genes like GSTs in muscadine grapes and will provide new research avenues to elucidate the mechanisms involved in flavonoid pathway during fruit development and ripening.

## Acknowledgements

We thank University of Arkansas System Division of Agriculture Fruit Research Station staff who maintained the plants and populations used in this study. We also thank Cindi Brownmiller for assistance with anthocyanin and proanthocyanidin quantification. Sequencing was performed by the University of Arkansas for Medical Sciences (UAMS). Funding was provided by the Hatch project ARK02599 and the NSF Student Undergraduate Research Fellowship (SURF) program.

## Authors Contributions

A.V. and M.W. provided overall conceptual guidance for the project and wrote the manuscript. A.V. and L.N. performed DNA and RNA extractions, cDNA synthesis, PCR, and gene expression analyses. A.V., M.W., L.N., and A.B. collected leaf and fruit samples used in this study. A.B. performed anthocyanin and proanthocyanidin analyses under the supervision and guidance of A.V., M.W., R.T., and L.H. J.R.C developed the mapping populations.

---

**Table S1:** BLASTn results from the sequence alignment of the 395 bp PCR product from genomic DNA of ‘Fry’ (Query1) and ‘Supreme’ (Query2) muscadines with *VaGST4* (Subject1) and VvGST4 (Subject2) sequences from *Vitis amurensis* and *V. vinifera*, respectively.

| Alignment | Identities (%) | Gaps (%) | Expect Value | Score (bits) |
| --- | --- | --- | --- | --- |
| Query1/Subject1 | 97.72 | 0 | 0.0 | 680 |
| Query1/Subject2 | 97.95 | 0 | 0.0 | 678 |
| Query2/Subject1 | 98.23 | 0 | 0.0 | 691 |
| Query2/Subject2 | 98.47 | 0 | 0.0 | 689 |

**Table S2.** Berry color phenotypes and KASP predicted *VrGST4* genotypes of 14 cultivars, 65 breeding selections, and 320 progeny from two muscadine mapping populations

| Population | Genotype ID | KASP<br>*VrGST4*<br>Genotype<sup>a</sup> | Phenotype | Full gene<br>sequencing<br>*VrGST4*<br>Genotype<sup>b</sup> |
| --- | --- | --- | --- | --- |
| Supreme x Nesbitt | 16-1 | C:C | black | na |
| Supreme x Nesbitt | 16-2 | T:T | bronze | na |
| Supreme x Nesbitt | 16-3 | C:T | black | na |
| Supreme x Nesbitt | 16-4 | C:C | black | na |
| Supreme x Nesbitt | 16-8 | C:T | black | na |
| Supreme x Nesbitt | 16-9 | C:C | black | na |
| Supreme x Nesbitt | 16-10 | T:T | bronze | na |
| Supreme x Nesbitt | 16-11 | C:T | black | na |
| Supreme x Nesbitt | 16-12 | C:T | bronze | C:T |
| Supreme x Nesbitt | 16-14 | na | black | na |
| Supreme x Nesbitt | 16-15 | na | bronze | na |
| Supreme x Nesbitt | 16-16 | C:T | black | na |
| Supreme x Nesbitt | 16-17 | C:T | black | na |
| Supreme x Nesbitt | 16-18 | C:C | black | na |
| Supreme x Nesbitt | 16-19 | C:C | black | na |
| Supreme x Nesbitt | 16-20 | C:C | black | na |
| Supreme x Nesbitt | 16-21 | C:T | black | na |
| Supreme x Nesbitt | 16-23 | C:T | black | na |
| Supreme x Nesbitt | 16-24 | T:T | bronze | na |
| Supreme x Nesbitt | 16-25 | C:C | black | na |
| Supreme x Nesbitt | 16-26 | C:C | black | na |
| Supreme x Nesbitt | 16-27 | T:T | bronze | na |
| Supreme x Nesbitt | 16-28 | C:C | black | na |
| Supreme x Nesbitt | 16-29 | T:T | bronze | na |
| Supreme x Nesbitt | 16-30 | C:T | black | na |
| Supreme x Nesbitt | 16-31 | T:T | bronze | na |
| Supreme x Nesbitt | 16-32 | T:T | bronze | na |
| Supreme x Nesbitt | 16-33 | C:T | black | na |
| Supreme x Nesbitt | 16-34 | T:T | bronze | na |
| Supreme x Nesbitt | 16-35 | C:T | black | na |
| Supreme x Nesbitt | 16-36 | C:T | black | na |
| Supreme x Nesbitt | 16-37 | C:T | black | na |
| Supreme x Nesbitt | 16-39 | C:T | black | na |
| Supreme x Nesbitt | 16-40 | C:C | black | na |
| Supreme x Nesbitt | 16-41 | C:T | black | na |
| Supreme x Nesbitt | 16-42 | T:T | bronze | na |
| Supreme x Nesbitt | 16-43 | T:T | bronze | na |
| Supreme x Nesbitt | 16-44 | C:C | black | na |
| Supreme x Nesbitt | 16-45 | C:T | black | na |
| Supreme x Nesbitt | 16-46 | T:T | bronze | na |
| Supreme x Nesbitt | 16-47 | C:T | black | na |
| Supreme x Nesbitt | 16-48 | T:T | bronze | na |
| Supreme x Nesbitt | 16-49 | C:T | black | na |
| Supreme x Nesbitt | 16-50 | C:C | black | na |
| Supreme x Nesbitt | 16-51 | C:T | black | na |
| Supreme x Nesbitt | 16-52 | C:C | black | na |
| Supreme x Nesbitt | 16-53 | C:T | black | na |
| Supreme x Nesbitt | 16-54 | C:T | -- | na |
| Supreme x Nesbitt | 16-55 | T:T | bronze | na |
| Supreme x Nesbitt | 16-56 | C:T | black | na |
| Supreme x Nesbitt | 16-57 | C:T | black | na |
| Supreme x Nesbitt | 16-58 | C:T | black | na |
| Supreme x Nesbitt | 16-59 | T:T | bronze | na |
| Supreme x Nesbitt | 16-60 | C:T | black | na |
| Supreme x Nesbitt | 16-61 | C:C | black | na |
| Supreme x Nesbitt | 16-62 | C:C | black | na |
| Supreme x Nesbitt | 16-63 | C:C | black | na |
| Supreme x Nesbitt | 16-64 | C:C | black | na |
| Supreme x Nesbitt | 16-65 | C:T | black | na |
| Supreme x Nesbitt | 16-66 | C:T | black | na |
| Supreme x Nesbitt | 16-67 | C:C | black | na |
| Supreme x Nesbitt | 16-68 | C:T | black | na |
| Supreme x Nesbitt | 16-69 | C:C | black | na |
| Supreme x Nesbitt | 16-70 | C:T | black | na |
| Supreme x Nesbitt | 16-71 | C:C | black | na |
| Supreme x Nesbitt | 16-72 | C:T | black | na |
| Supreme x Nesbitt | 16-73 | C:T | black | na |
| Supreme x Nesbitt | 16-74 | C:C | black | na |
| Supreme x Nesbitt | 16-75 | C:T | black | na |
| Supreme x Nesbitt | 16-76 | C:T | black | na |
| Supreme x Nesbitt | 16-77 | C:C | black | na |
| Supreme x Nesbitt | 16-78 | C:T | black | na |
| Supreme x Nesbitt | 16-79 | C:T | black | na |
| Supreme x Nesbitt | 16-80 | C:C | black | na |
| Supreme x Nesbitt | 16-81 | C:T | black | na |
| Supreme x Nesbitt | 16-82 | C:T | black | na |
| Supreme x Nesbitt | 16-83 | C:C | black | na |
| Supreme x Nesbitt | 16-84 | C:T | black | na |
| Supreme x Nesbitt | 16-85 | C:T | black | na |
| Supreme x Nesbitt | 16-86 | C:T | black | na |
| Supreme x Nesbitt | 16-87 | C:C | black | na |
| Supreme x Nesbitt | 16-88 | C:C | bronze | T:T |
| Supreme x Nesbitt | 16-89 | T:T | black | C:T |
| Supreme x Nesbitt | 16-90 | C:T | black | na |
| Supreme x Nesbitt | 16-91 | C:T | black | na |
| Supreme x Nesbitt | 16-92 | C:T | black | na |
| Supreme x Nesbitt | 16-93 | T:T | bronze | na |
| Supreme x Nesbitt | 16-94 | C:C | black | na |
| Supreme x Nesbitt | 16-95 | na | black | na |
| Supreme x Nesbitt | 16-96 | C:C | black | na |
| Supreme x Nesbitt | 16-97 | C:C | black | na |
| Supreme x Nesbitt | 16-101 | C:C | black | na |
| Supreme x Nesbitt | 16-102 | C:C | black | na |
| Supreme x Nesbitt | 16-103 | C:T | black | na |
| Supreme x Nesbitt | 16-104 | C:T | black | na |
| Supreme x Nesbitt | 16-105 | T:T | bronze | na |
| Supreme x Nesbitt | 16-106 | C:T | black | na |
| Supreme x Nesbitt | 16-107 | C:T | black | na |
| Supreme x Nesbitt | 16-109 | C:T | black | na |
| Supreme x Nesbitt | 16-110 | C:C | black | na |
| Supreme x Nesbitt | 16-111 | T:T | bronze | na |
| Supreme x Nesbitt | 16-112 | na | bronze | na |
| Supreme x Nesbitt | 16-114 | C:C | black | na |
| Supreme x Nesbitt | 16-115 | C:C | black | na |
| Supreme x Nesbitt | 16-116 | C:C | black | na |
| Supreme x Nesbitt | 16-117 | C:T | black | na |
| Supreme x Nesbitt | 16-118 | C:C | black | na |
| Supreme x Nesbitt | 16-119 | C:T | black | na |
| Supreme x Nesbitt | 16-120 | C:T | black | na |
| Supreme x Nesbitt | 16-121 | C:T | black | na |
| Supreme x Nesbitt | 16-122 | C:C | black | na |
| Supreme x Nesbitt | 16-123 | C:T | black | na |
| Supreme x Nesbitt | 16-124 | C:T | black | na |
| Supreme x Nesbitt | 16-125 | C:T | bronze | T:T |
| Supreme x Nesbitt | 16-126 | T:T | bronze | na |
| Supreme x Nesbitt | 16-127 | na | black | na |
| Supreme x Nesbitt | 16-128 | T:T | bronze | na |
| Supreme x Nesbitt | 16-129 | T:T | bronze | na |
| Supreme x Nesbitt | 16-130 | C:C | black | na |
| Supreme x Nesbitt | 16-131 | C:T | black | na |
| Supreme x Nesbitt | 16-132 | T:T | bronze | na |
| Supreme x Nesbitt | 16-133 | C:T | black | na |
| Supreme x Nesbitt | 16-134 | na | black | na |
| Supreme x Nesbitt | 16-135 | C:C | black | na |
| Supreme x Nesbitt | 16-136 | C:T | black | na |
| Supreme x Nesbitt | 16-137 | C:T | black | na |
| Supreme x Nesbitt | 16-138 | C:T | black | na |
| Supreme x Nesbitt | 16-139 | na | black | na |
| Supreme x Nesbitt | 16-140 | C:C | black | na |
| Supreme x Nesbitt | 16-141 | T:T | bronze | na |
| Supreme x Nesbitt | 16-142 | C:T | black | na |
| Supreme x Nesbitt | 16-143 | T:T | bronze | na |
| Supreme x Nesbitt | 16-144 | T:T | bronze | na |
| Supreme x Nesbitt | 16-145 | C:T | black | na |
| Supreme x Nesbitt | 16-146 | C:T | black | na |
| Supreme x Nesbitt | 16-147 | C:T | black | na |
| Supreme x Nesbitt | 16-148 | C:T | black | na |
| Supreme x Nesbitt | 16-149 | C:T | black | na |
| Supreme x Nesbitt | 16-150 | T:T | bronze | na |
| Supreme x Nesbitt | 16-151 | C:C | black | na |
| Supreme x Nesbitt | 16-152 | C:C | black | na |
| Supreme x Nesbitt | 16-153 | C:T | black | na |
| Supreme x Nesbitt | 16-154 | C:T | black | na |
| Supreme x Nesbitt | 16-155 | C:C | black | na |
| Supreme x Nesbitt | 16-156 | C:C | black | na |
| Supreme x Nesbitt | 16-157 | C:T | black | na |
| Supreme x Nesbitt | 16-158 | C:T | black | na |
| Supreme x Nesbitt | 16-159 | T:T | bronze | na |
| Supreme x Nesbitt | 16-160 | C:C | black | na |
| Supreme x Nesbitt | 16-161 | C:C | black | na |
| Supreme x Nesbitt | 16-162 | C:T | black | na |
| Supreme x Nesbitt | 16-163 | C:T | black | na |
| Supreme x Nesbitt | 16-164 | na | black | na |
| Supreme x Nesbitt | 16-165 | C:T | black | na |
| Supreme x Nesbitt | 16-166 | C:T | black | na |
| Supreme x Nesbitt | 16-167 | C:T | black | na |
| Supreme x Nesbitt | 16-168 | C:T | black | na |
| Supreme x Nesbitt | 16-169 | C:C | black | na |
| Supreme x Nesbitt | 16-170 | C:C | black | na |
| Supreme x Nesbitt | 16-171 | C:C | black | na |
| Supreme x Nesbitt | 16-172 | T:T | bronze | na |
| Supreme x Nesbitt | 16-173 | C:C | black | na |
| Supreme x Nesbitt | 16-174 | C:T | black | na |
| Black Beauty x Nesbitt | 17-1 | C:T | black | na |
| Black Beauty x Nesbitt | 17-2 | C:T | black | na |
| Black Beauty x Nesbitt | 17-3 | C:T | black | na |
| Black Beauty x Nesbitt | 17-4 | C:T | black | na |
| Black Beauty x Nesbitt | 17-5 | C:C | black | na |
| Black Beauty x Nesbitt | 17-6 | C:T | black | na |
| Black Beauty x Nesbitt | 17-7 | C:T | black | na |
| Black Beauty x Nesbitt | 17-8 | C:C | black | na |
| Black Beauty x Nesbitt | 17-9 | C:C | -- | na |
| Black Beauty x Nesbitt | 17-10 | T:T | bronze | na |
| Black Beauty x Nesbitt | 17-11 | T:T | bronze | na |
| Black Beauty x Nesbitt | 17-12 | C:C | black | na |
| Black Beauty x Nesbitt | 17-13 | C:T | -- | na |
| Black Beauty x Nesbitt | 17-14 | C:T | black | na |
| Black Beauty x Nesbitt | 17-15 | C:T | black | na |
| Black Beauty x Nesbitt | 17-16 | C:T | -- | na |
| Black Beauty x Nesbitt | 17-17 | C:T | black | na |
| Black Beauty x Nesbitt | 17-20 | T:T | bronze | na |
| Black Beauty x Nesbitt | 17-21 | C:T | black | na |
| Black Beauty x Nesbitt | 17-22 | C:T | black | na |
| Black Beauty x Nesbitt | 17-23 | C:C | black | na |
| Black Beauty x Nesbitt | 17-24 | T:T | bronze | na |
| Black Beauty x Nesbitt | 17-25 | C:C | black | na |
| Black Beauty x Nesbitt | 17-26 | C:T | black | na |
| Black Beauty x Nesbitt | 17-27 | C:T | black | na |
| Black Beauty x Nesbitt | 17-28 | C:C | -- | na |
| Black Beauty x Nesbitt | 17-29 | C:T | -- | na |
| Black Beauty x Nesbitt | 17-30 | T:T | bronze | na |
| Black Beauty x Nesbitt | 17-32 | C:C | black | na |
| Black Beauty x Nesbitt | 17-33 | na | bronze | na |
| Black Beauty x Nesbitt | 17-34 | na | bronze | na |
| Black Beauty x Nesbitt | 17-35 | C:C | black | na |
| Black Beauty x Nesbitt | 17-36 | C:C | black | na |
| Black Beauty x Nesbitt | 17-37 | C:T | black | na |
| Black Beauty x Nesbitt | 17-38 | C:T | black | na |
| Black Beauty x Nesbitt | 17-39 | C:C | black | na |
| Black Beauty x Nesbitt | 17-40 | C:C | black | na |
| Black Beauty x Nesbitt | 17-42 | T:T | bronze | na |
| Black Beauty x Nesbitt | 17-43 | C:T | black | na |
| Black Beauty x Nesbitt | 17-44 | C:T | black | na |
| Black Beauty x Nesbitt | 17-45 | C:T | black | na |
| Black Beauty x Nesbitt | 17-46 | C:T | black | na |
| Black Beauty x Nesbitt | 17-47 | C:C | black | na |
| Black Beauty x Nesbitt | 17-50 | C:T | black | na |
| Black Beauty x Nesbitt | 17-52 | C:T | -- | na |
| Black Beauty x Nesbitt | 17-53 | C:T | black | na |
| Black Beauty x Nesbitt | 17-54 | C:T | black | na |
| Black Beauty x Nesbitt | 17-55 | T:T | -- | na |
| Black Beauty x Nesbitt | 17-56 | C:T | -- | na |
| Black Beauty x Nesbitt | 17-57 | C:T | -- | na |
| Black Beauty x Nesbitt | 17-58 | C:T | black | na |
| Black Beauty x Nesbitt | 17-59 | C:T | -- | na |
| Black Beauty x Nesbitt | 17-60 | T:T | bronze | na |
| Black Beauty x Nesbitt | 17-61 | C:T | black | na |
| Black Beauty x Nesbitt | 17-62 | C:T | black | na |
| Black Beauty x Nesbitt | 17-64 | C:T | black | na |
| Black Beauty x Nesbitt | 17-65 | T:T | bronze | na |
| Black Beauty x Nesbitt | 17-66 | C:T | black | na |
| Black Beauty x Nesbitt | 17-68 | T:T | bronze | na |
| Black Beauty x Nesbitt | 17-69 | T:T | bronze | na |
| Black Beauty x Nesbitt | 17-70 | C:C | black | na |
| Black Beauty x Nesbitt | 17-71 | T:T | -- | na |
| Black Beauty x Nesbitt | 17-72 | T:T | bronze | na |
| Black Beauty x Nesbitt | 17-73 | C:C | black | na |
| Black Beauty x Nesbitt | 17-76 | C:T | black | na |
| Black Beauty x Nesbitt | 17-77 | C:C | black | na |
| Black Beauty x Nesbitt | 17-78 | C:C | black | na |
| Black Beauty x Nesbitt | 17-79 | C:T | black | na |
| Black Beauty x Nesbitt | 17-80 | T:T | bronze | na |
| Black Beauty x Nesbitt | 17-81 | T:T | bronze | na |
| Black Beauty x Nesbitt | 17-82 | T:T | bronze | na |
| Black Beauty x Nesbitt | 17-83 | C:T | black | na |
| Black Beauty x Nesbitt | 17-84 | C:T | black | na |
| Black Beauty x Nesbitt | 17-85 | C:T | black | na |
| Black Beauty x Nesbitt | 17-86 | C:T | black | na |
| Black Beauty x Nesbitt | 17-87 | C:T | black | na |
| Black Beauty x Nesbitt | 17-88 | T:T | bronze | na |
| Black Beauty x Nesbitt | 17-89 | C:T | black | na |
| Black Beauty x Nesbitt | 17-90 | C:T | black | na |
| Black Beauty x Nesbitt | 17-91 | C:T | -- | na |
| Black Beauty x Nesbitt | 17-92 | T:T | bronze | na |
| Black Beauty x Nesbitt | 17-93 | C:T | black | na |
| Black Beauty x Nesbitt | 17-94 | C:T | black | na |
| Black Beauty x Nesbitt | 17-95 | C:T | black | na |
| Black Beauty x Nesbitt | 17-96 | C:C | black | na |
| Black Beauty x Nesbitt | 17-97 | C:T | -- | na |
| Black Beauty x Nesbitt | 17-98 | C:T | black | na |
| Black Beauty x Nesbitt | 17-99 | T:T | bronze | na |
| Black Beauty x Nesbitt | 17-100 | C:T | black | na |
| Black Beauty x Nesbitt | 17-101 | C:T | black | na |
| Black Beauty x Nesbitt | 17-102 | na | -- | na |
| Black Beauty x Nesbitt | 17-103 | C:C | black | na |
| Black Beauty x Nesbitt | 17-104 | C:T | black | na |
| Black Beauty x Nesbitt | 17-105 | C:T | black | na |
| Black Beauty x Nesbitt | 17-106 | C:T | black | na |
| Black Beauty x Nesbitt | 17-107 | C:T | black | na |
| Black Beauty x Nesbitt | 17-108 | C:T | black | na |
| Black Beauty x Nesbitt | 17-109 | T:T | bronze | na |
| Black Beauty x Nesbitt | 17-110 | C:T | black | na |
| Black Beauty x Nesbitt | 17-111 | T:T | bronze | na |
| Black Beauty x Nesbitt | 17-112 | T:T | bronze | na |
| Black Beauty x Nesbitt | 17-113 | C:C | black | na |
| Black Beauty x Nesbitt | 17-114 | na | black | na |
| Black Beauty x Nesbitt | 17-115 | C:T | black | na |
| Black Beauty x Nesbitt | 17-116 | C:C | black | na |
| Black Beauty x Nesbitt | 17-117 | C:T | black | na |
| Black Beauty x Nesbitt | 17-118 | T:T | bronze | na |
| Black Beauty x Nesbitt | 17-119 | C:T | black | na |
| Black Beauty x Nesbitt | 17-120 | C:T | black | na |
| Black Beauty x Nesbitt | 17-121 | C:C | black | na |
| Black Beauty x Nesbitt | 17-122 | C:C | black | na |
| Black Beauty x Nesbitt | 17-123 | C:C | black | na |
| Black Beauty x Nesbitt | 17-124 | C:T | -- | na |
| Black Beauty x Nesbitt | 17-125 | C:T | black | na |
| Black Beauty x Nesbitt | 17-126 | C:T | black | na |
| Black Beauty x Nesbitt | 17-129 | T:T | bronze | na |
| Black Beauty x Nesbitt | 17-130 | C:T | black | na |
| Black Beauty x Nesbitt | 17-131 | C:T | black | na |
| Black Beauty x Nesbitt | 17-132 | C:C | black | na |
| Black Beauty x Nesbitt | 17-133 | C:T | black | na |
| Black Beauty x Nesbitt | 17-134 | T:T | bronze | na |
| Black Beauty x Nesbitt | 17-135 | T:T | bronze | na |
| Black Beauty x Nesbitt | 17-136 | C:C | black | na |
| Black Beauty x Nesbitt | 17-137 | T:T | bronze | na |
| Black Beauty x Nesbitt | 17-138 | C:C | black | na |
| Black Beauty x Nesbitt | 17-139 | C:T | black | na |
| Black Beauty x Nesbitt | 17-140 | T:T | bronze | na |
| Black Beauty x Nesbitt | 17-141 | T:T | bronze | na |
| Black Beauty x Nesbitt | 17-142 | C:T | black | na |
| Black Beauty x Nesbitt | 17-143 | T:T | bronze | na |
| Black Beauty x Nesbitt | 17-144 | C:T | black | na |
| Black Beauty x Nesbitt | 17-145 | C:T | black | na |
| Black Beauty x Nesbitt | 17-146 | C:C | black | na |
| Black Beauty x Nesbitt | 17-148 | C:T | black | na |
| Black Beauty x Nesbitt | 17-149 | C:T | black | na |
| Black Beauty x Nesbitt | 17-150 | C:T | black | na |
| Black Beauty x Nesbitt | 17-151 | C:T | black | na |
| Black Beauty x Nesbitt | 17-152 | C:T | black | na |
| Black Beauty x Nesbitt | 17-153 | na | black | na |
| Black Beauty x Nesbitt | 17-154 | T:T | bronze | na |
| Black Beauty x Nesbitt | 17-155 | na | black | na |
| Black Beauty x Nesbitt | 17-156 | C:C | black | na |
| Black Beauty x Nesbitt | 17-157 | C:T | black | na |
| Black Beauty x Nesbitt | 17-158 | C:T | black | na |
| Black Beauty x Nesbitt | 17-159 | C:T | black | na |
| Black Beauty x Nesbitt | 17-160 | C:T | black | na |
| Black Beauty x Nesbitt | 17-161 | T:T | bronze | na |
| Black Beauty x Nesbitt | 17-162 | C:C | black | na |
| Black Beauty x Nesbitt | 17-163 | T:T | bronze | na |
| Black Beauty x Nesbitt | 17-164 | C:T | black | na |
| Black Beauty x Nesbitt | 17-165 | T:T | bronze | na |
| Black Beauty x Nesbitt | 17-166 | T:T | bronze | na |
| Black Beauty x Nesbitt | 17-167 | C:T | black | na |
| Black Beauty x Nesbitt | 17-168 | T:T | bronze | na |
| Black Beauty x Nesbitt | 17-169 | C:T | black | na |
| Black Beauty x Nesbitt | 17-171 | C:T | black | na |
| Black Beauty x Nesbitt | 17-172 | na | bronze | na |
| Breeding Selection | AM-9 | C:T | black | na |
| Breeding Selection | AM-26 | T:T | bronze | na |
| Breeding Selection | AM-27 | C:T | black | na |
| Breeding Selection | AM-41 | C:T | black | na |
| Breeding Selection | AM-45 | T:T | bronze | na |
| Breeding Selection | AM-53 | T:T | bronze | na |
| Breeding Selection | AM-54 | T:T | bronze | na |
| Breeding Selection | AM-64 | C:T | black | na |
| Breeding Selection | AM-70 | C:T | black | na |
| Breeding Selection | AM-71 | T:T | bronze | na |
| Breeding Selection | AM-77 | C:T | black | na |
| Breeding Selection | AM-81 | T:T | bronze | na |
| Breeding Selection | AM-83 | C:T | black | na |
| Breeding Selection | AM-87 | C:T | black | na |
| Breeding Selection | AM-90 | T:T | bronze | na |
| Breeding Selection | AM-92 | T:T | bronze | na |
| Breeding Selection | AM-102 | C:T | black | na |
| Breeding Selection | AM-105 | T:T | bronze | na |
| Breeding Selection | AM-120 | T:T | bronze | na |
| Breeding Selection | AM-121 | C:T | black | na |
| Breeding Selection | AM-126 | T:T | bronze | na |
| Breeding Selection | AM-127 | C:C | black | na |
| Breeding Selection | AM-131 | T:T | bronze | na |
| Breeding Selection | AM-132 | C:T | black | na |
| Breeding Selection | AM-135 | T:T | bronze | na |
| Breeding Selection | AM-136 | C:T | black | na |
| Breeding Selection | AM-137 | C:T | black | na |
| Breeding Selection | AM-142 | C:T | black | na |
| Breeding Selection | AM-143 | C:T | black | na |
| Breeding Selection | AM-148 | C:T | black | na |
| Breeding Selection | AM-150 | C:T | black | na |
| Breeding Selection | AM-156 | T:T | bronze | na |
| Breeding Selection | AM-158 | T:T | bronze | na |
| Breeding Selection | AM-161 | C:T | black | na |
| Breeding Selection | AM-165 | T:T | bronze | na |
| Breeding Selection | AM-167 | T:T | bronze | na |
| Breeding Selection | AM-169 | C:T | black | na |
| Breeding Selection | AM-170 | T:T | bronze | na |
| Breeding Selection | AM-171 | na | black | na |
| Breeding Selection | AM-172 | C:T | black | na |
| Breeding Selection | AM-173 | T:T | bronze | na |
| Breeding Selection | AM-174 | C:T | black | na |
| Breeding Selection | AM-175 | T:T | black | C:T |
| Breeding Selection | AM-176 | T:T | bronze | na |
| Breeding Selection | AM-177 | na | black | na |
| Breeding Selection | AM-178 | C:T | black | na |
| Breeding Selection | AM-179 | T:T | bronze | na |
| Breeding Selection | AM-181 | C:T | black | na |
| Breeding Selection | AM-182 | C:T | black | na |
| Breeding Selection | AM-183 | C:T | black | na |
| Breeding Selection | AM-184 | C:T | black | na |
| Breeding Selection | AM-185 | C:T | black | na |
| Breeding Selection | AM-186 | T:T | bronze | na |
| Breeding Selection | AM-187 | C:T | black | na |
| Breeding Selection | AM-188 | C:T | black | na |
| Breeding Selection | AM-189 | C:T | black | na |
| Breeding Selection | AM-190 | C:T | black | na |
| Breeding Selection | AM-191 | C:C | bronze | T:T |
| Breeding Selection | AM-192 | C:T | black | na |
| Breeding Selection | AM-193 | T:T | bronze | na |
| Breeding Selection | AM-194 | C:T | black | na |
| Breeding Selection | AM-195 | C:T | black | na |
| Breeding Selection | AM-196 | C:T | black | na |
| Breeding Selection | AM-197 | C:T | black | na |
| Breeding Selection | AM-198 | C:T | black | na |
| Cultivar | Black Beauty | C:T | black | na |
| Cultivar | Carlos | T:T | bronze | na |
| Cultivar | Delicious | C:T | black | na |
| Cultivar | Doreen | T:T | bronze | na |
| Cultivar | Fry | T:T | bronze | na |
| Cultivar | Granny Val | T:T | bronze | na |
| Cultivar | Ison | C:T | black | na |
| Cultivar | Nesbitt | C:T | black | na |
| Cultivar | Noble | C:C | black | na |
| Cultivar | Southern Home | C:T | black | na |
| Cultivar | Sugargate | C:T | black | na |
| Cultivar | Summit | T:T | bronze | na |
| Cultivar | Supreme | C:T | black | na |
| Cultivar | Tara | T:T | bronze | na |

<sup>a</sup> C:C = homozygote black, C:T = heterozygote black, T:T = bronze, na = failed reaction or
<sup>b</sup> The full *VrGST4* gene sequence was obtained for genotypes 16-12, 16-88, 16-89, 16-125, AM-175, and AM-191 because of discrepancies between the KASP marker data and phenotypes

**Figure S1.**
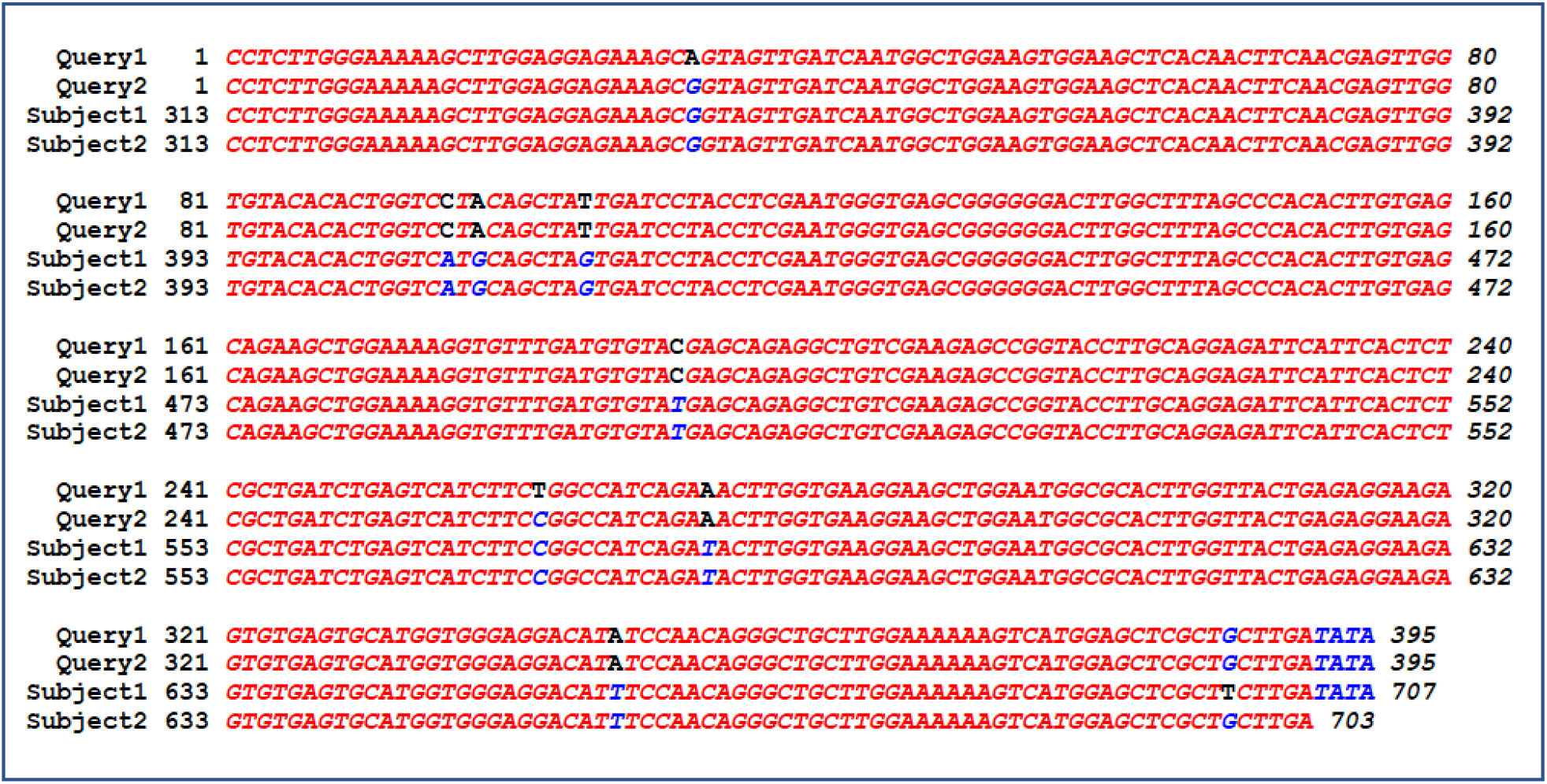
Sequence alignment of the 395 bp PCR product from genomic DNA of ‘Fry’ (Query1) and ‘Supreme’ (Query2) muscadines with *VaGST4* (Subject1) and *VvGST4* (Subject2) sequences from *Vitis amurensis* and *V. vinifera*, respectively. Numbers in the alignment represent base pairs.

